# Higher order phosphatase-substrate contacts terminate the Integrated Stress Response

**DOI:** 10.1101/2021.06.18.449003

**Authors:** Yahui Yan, Heather P. Harding, David Ron

## Abstract

Many regulatory PPP1R subunits join few catalytic PP1c subunits to mediate phosphoserine and phosphothreonine dephosphorylation in metazoans. Regulatory subunits are known to engage PP1c’s surface, locally affecting flexible phosphopeptides access to the active site. However, catalytic efficiency of holophosphatases towards their natively-folded phosphoprotein substrates is largely unexplained. Here we present a Cryo-EM structure of the tripartite PP1c/PPP1R15A/G-actin holophosphatase that terminates signalling in the Integrated Stress Response (ISR) in pre-dephosphorylation complex with its substrate, translation initiation factor 2α (eIF2α). G-actin’s role in eIF2α dephosphorylation is supported crystallographically by the structure of the binary PPP1R15A-G-actin complex, and by biochemical and genetic confirmation of the essential role of PPP1R15A-G-actin contacts to eIF2α^P^ dephosphorylation. In the pre-dephosphorylation CryoEM complex, G-actin aligns the catalytic and regulatory subunits, creating a composite surface that engages eIF2α’s N-terminal domain to position the distant phosphoserine-51 at the active site. eIF2α residues specifying affinity for the holophosphatase are confirmed here to make critical contacts with the eIF2α kinase PERK. Thus, a convergent process of higher-order substrate recognition specifies functionally-antagonistic phosphorylation and dephosphorylation in the ISR.

## INTRODUCTION

Signalling in many pathways is terminated by dephosphorylation of pSer and pThr. This task is relegated to complexes between a small number of related protein phosphatase catalytic subunits and hundreds of different regulatory subunits that impart substrate specificity on the holophosphatases (Brautigan, 2013). A diverse group of regulatory subunits bind members of the PP1 family of related catalytic subunits (Heroes *et al*, 2013; Verbinnen *et al*, 2017).

The structural basis of complex formation between the PPP1R, regulatory, and PP1c, catalytic subunits, is understood. The former engage the surface of PP1c via short linear segments, exemplified by the conserved RVxF, *ϕϕ* and ‘R’ motives observed in complexes of G_M_ (Egloff *et al*, 1997), spinophilin (Ragusa *et al*, 2010), PNUTS (Choy *et al*, 2014) or Phactr1 (Fedoryshchak *et al*, 2020) regulatory subunits with PP1c. Regulatory subunit binding sculpts the surface of PP1c to bias substrate access to the active site. This explains local selectivity for and against flexible pSer/pThr-bearing substrate-derived peptides (Heroes *et al*., 2013; Li *et al*, 2013; Peti *et al*, 2013; Roy & Cyert, 2009), but cannot fully account for the catalytic efficiency of holophosphatases directed towards their physiological substrates - globular domains of phosphoproteins.

Eukaryotes share a signal transduction pathway that couples changing rates of translation initiation to a pervasive transcriptional and translational programme. Known in yeasts as the General Control Response (Hinnebusch, 2005) and in animals as the Integrated Stress Response (ISR) (Harding *et al*, 2003), the ISR is triggered by phosphorylation of Ser51 of the α-subunit of translation initiation factor 2 (eIF2α^P^), an event mediated by four known kinases (GCN2, PERK, HRI and PKR). ISR manipulation reveals its broad role in cellular homeostasis and the potential for tuning the response to therapeutic ends (Costa-Mattioli & Walter, 2020).

Dephosphorylation of eIF2α^P^ terminates signalling in the ISR. In mammals dephosphorylation is assisted by one of two known regulatory subunits, PPP1R15A or B. PPP1R15A, also known as GADD34, is encoded by an ISR-inducible gene and serves in negative feedback (Brush *et al*, 2003; Ma & Hendershot, 2003; Novoa *et al*, 2001; Novoa *et al*, 2003) whereas PPP1R15B, also known as CReP, is constitutively expressed (Jousse *et al*, 2003). Genetic studies demonstrate the benefits of extending the ISR and suggest the therapeutic potential of inhibiting eIF2α dephosphorylation (D’Antonio *et al*, 2013; Marciniak *et al*, 2004). Attaining this goal requires a detailed understanding of the enzyme(s) involved.

The mammalian PPP1R15s are proteins of more than 600 amino acids, but their conserved portion is limited to ∼70-residues of their C-termini. This segment (residues 555-621 in human PPP1R15A), which is sufficient to promote eIF2α^P^ dephosphorylation when expressed in cells (Jousse *et al*., 2003; Novoa *et al*., 2001) is also conserved in the single PPP1R15 gene of other animal phyla (Malzer *et al*, 2013) and in a pathogenicity gene of Herpes viruses (He *et al*, 1997). The N-terminal half of this conserved PPP1R15 core binds PP1c (Chen *et al*, 2015; Choy *et al*, 2015; Connor *et al*, 2001; Novoa *et al*., 2001). However, the PP1c/PPP1R15 complex is little better at dephosphorylating eIF2α^P^ than a PP1c apo-enzyme (Chen *et al*., 2015; Crespillo-Casado *et al*, 2018). High resolution crystal structures of PP1c in complex with PPP1R15A (Choy *et al*., 2015) and PPP1R15B (Chen *et al*., 2015) exist, however, these have merely generic features of PP1c holoenzymes. Addition of G-actin strongly stimulates eIF2α^P^ dephosphorylation by PP1c/PPP1R15. G-actin binds the C-terminal half of the conserved core of the PPP1R15s to form a ternary PP1c/PPP1R15/G-actin complex both in vitro (Chen *et al*., 2015; Crespillo-Casado *et al*, 2017) and in cell extracts (Chambers *et al*, 2015), but the basis of its stimulatory activity remains unknown.

Here we report on a crystal structure of the binary PPP1R15A/G-actin complex and on a CryoEM structure of the tripartite PP1c/PPP1R15A/G-actin holophosphatase in complex with its substrate. Comparison of this eIF2α^P^ pre-dephosphorylation complex with a counterpart eIF2α pre-phosphorylation complex (with the kinase PKR (Dar *et al*, 2005)) reveals similar principles by which a protein kinase and a PP1-based holophosphatases attain substrate-specific catalytic efficiency.

## RESULTS

### Crystal structure of the binary PPP1R15A/G-actin complex

Enzymatic and binding experiments suggested that the conserved core of PPP1R15s binds PP1c and G-actin via non-overlapping linear segments (Fig. 1A). High resolution crystal structures of binary PPP1R15A/PP1 and PPP1R15B/PP1 complexes exist (Chen *et al*., 2015; Choy *et al*., 2015), but similar information on contacts with G-actin is missing. We exploited the observation that G-actin can be recruited to the PPP1R15 containing holophosphatase in complex with DNase I (Chen *et al*., 2015). DNase I stabilises actin monomers (Mannherz *et al*, 1980) facilitates crystallisation (Kabsch *et al*, 1990). DNase I-G-actin formed a stable complex with the C-terminal portion of PPP1R15A, which crystallised and diffracted to 2.55Å (Fig. 1B – 1D, supplementary fig. 1A and Table 1).

**Figure 1.**
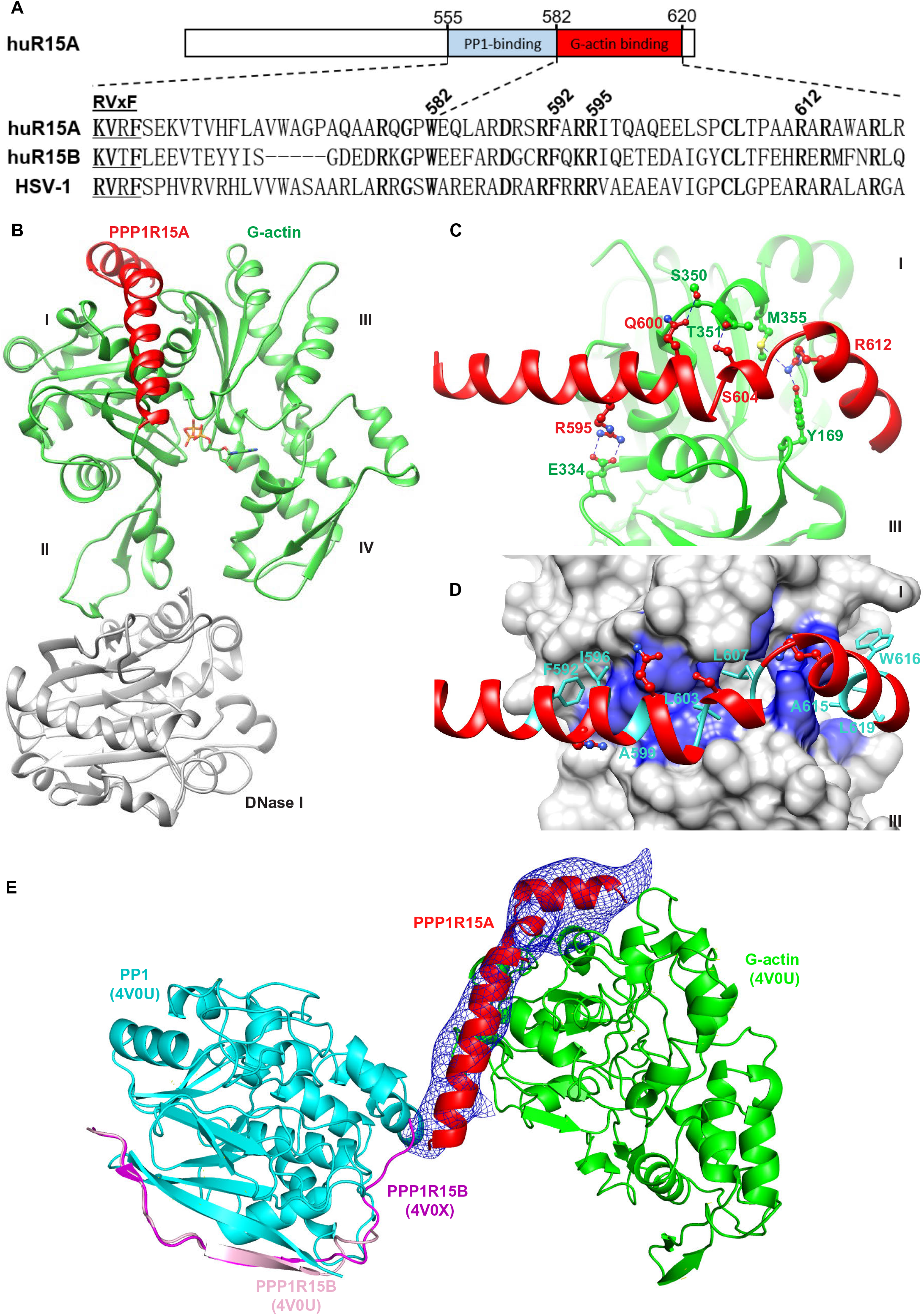
PPP1R15A engages G-actin’s barbed end. (A) Schema of human PPP1R15A. The PP1-and G-actin-binding segments are marked. Beneath is an alignment of human PPP1R15A (huR15A), human PPP1R15B (huR15B) and the regulatory subunit of Herpes simplex virus (HSV-1). Conserved residues bolded. The ‘RVxF’ motif, common to many PPP1Rs, is noted. Numbering refers to human PPP1R15A. (B) Ribbon diagram of the overall crystallographic structure of the PPP1R15A/G-actin/DNase I complex at 2.55 Å. Actin domains are numbered. (C) Close up view of hydrogen-bonding interactions between PPP1R15A (red) and G-actin (green). The density map for PPPR15A is shown in supplementary fig. 1A. (D) Hydrophobic interactions of G-actin (blue surface) with the indicated PPP1R15A residues (sidechains as turquoise sticks). (E) Model of a tripartite holophosphatase constructed by aligning the PPP1R15A/G-actin/DNase I complex (above) and the binary PP1G/PPP1R15B complex (PDB 4V0X) to the low resolution PP1G/PPP1R15B/G-actin complex (PDB 4V0U), the former via G-actin and latter via PP1. PP1G from PDB 4V0X (with RMSD of 0.318 Å between PP1c’s 290 Cα pairs) and G-actin from PPP1R15A/G-actin/DNase I complex (with RMSD of 0.626 Å between actin’s 343 Cα pairs) are not shown for clarity. Note the proximity of the C-terminus of PPP1R15B and the N-terminus of PPP1R15A (comprised of the same residue of both orthologues, PPP1R15B Trp662 & PPP1R15A Trp582) in the two binary complexes. The previously unaccounted density in G-actin’s barbed end (displayed as an average difference electron density from PDB 4V0U and shown as a blue mesh) accommodates the actin-binding helices of PPP1R15A from the aligned PPP1R15A/G-actin/DNase I complex.

**Table 1:** X-ray data collection and refinement statistics.

|  | **G-actin/DNase I**<br>**/PPP1R15A<sup>582-621</sup> complex** |
| --- | --- |
| **Data collection** |  |
| Synchrotron stations | DLS i03 |
| Space group | P2<sub>1</sub>2<sub>1</sub>2<sub>1</sub> |
| a,b,c (Å) | 86.48, 107.79, 192.42 |
| $\alpha, \beta, \gamma$ (°) | 90.00, 90.00, 90.00 |
| Resolution (Å) | 55.12-2.55 (2.62-2.55)<sup>a</sup> |
| Rmerge | 0.315 (1.701)<sup>a</sup> |
| $\langle I/\sigma(I) \rangle$ | 6.5 (1.4)<sup>a</sup> |
| CC1/2 | 0.987 (0.631)<sup>a</sup> |
| No. of unique reflections | 59509 (4573)<sup>a</sup> |
| Completeness, % | 100.0 (100.0)<sup>a</sup> |
| Redundancy | 6.9 (7.1)<sup>a</sup> |
| **Refinement** |  |
| Rwork/Rfree | 0.218/0.248 |
| No. of atoms (non H) | 10650 |
| Average B-factors | 36.1 |
| RMS Bond lengths (Å) | 0.002 |
| RMS Bond angles (°) | 1.209 |
| Ramachandran favoured region (%) | 97.9 |
| Ramachandran outliers (%) | 0 |
| MolProbity score | 0.78 |
| PDB code | 7NXV |

<sup>a</sup> Values in parentheses are for the highest resolution shell.

Residues 582-619 of PPP1R15A form two helices separated by a sharp turn that wrap around G-actin, engaging the groove between subdomains I and III. DNase I binds the opposite end of G-actin, whereas D-loop insertion in F-actin, and G-actin-binding by small molecules such as cytochalasin D overlap the PPP1R15A site (Supplementary fig. 1B & supplementary fig. 1C). These findings explain cytochalasin’s ability to antagonise the stimulatory effect of G-actin on eIF2α^P^ dephosphorylation in vitro (Chen *et al*., 2015) and the inhibitory effect of drug-enforced actin polymerisation on dephosphorylation in cells (Chambers *et al*., 2015).

PPP1R15A’s location on the surface of G-actin corresponds to a site of electron density observed in a crystal structure of the PP1G/PPP1R15B/G-actin complex (PDB 4V0U (Chen *et al*., 2015)). Due to low resolution of the map (7.8 Å), the density remained unassigned. However, aligning G-actin in the ternary PP1G/PPP1R15B/G-actin (PDB 4V0U) and binary PPP1R15A/G-actin/DNase I complex (here), reveals that the PPP1R15A from the high-resolution binary complex, fits nicely in the unassigned density in G-actin’s groove in PDB 4V0U. Furthermore, the C-terminal PPP1R15B Trp662 from a high resolution binary PP1G/PPP1R15B complex (PDB 4V0X, aligned by its PP1 to PDB 4V0U) is close enough to the N-terminal PPP1R15A orthologous residue (Trp582) in its complex with G-actin (here) to complete a PPP1R15 peptide chain in the ternary complex and provide a view of a composite PPP1R15B/A-containing tripartite holoenzyme (Fig. 1E).

### Validation of actin’s role as a co-factor of the eIF2α^P^ phosphatase in vitro and in cells

Residues lining one face of the PPP1R15A helical extension form hydrophobic interactions and hydrogen bonds with G-actin (Fig. 1C & 1D). Human PPP1R15A Phe592 inserts into a hydrophobic cavity on actin’s surface, whereas Arg595 forms a salt bridge with actin Asp334 and PPP1R15A Arg612 forms hydrogen bonds with actin Tyr169 and Met355. These PPP1R15A residues are conserved across species and were selected for functional studies by mutagenesis.

Recombinant wildtype or mutant human PPP1R15A core fragment (residues 553-624) in complex with PP1A and G-actin were compared in dephosphorylating the N-terminal lobe of eIF2α^P^. A kinetic defect was observed in all four mutants tested (Fig. 2A, 2B). The defect was selective for actin-containing mutant holophosphatases - the low baseline activity of the apo-enzymes (lacking G-actin) was barely affected by the mutations. The strongest defect was observed in double mutants compromising both PPP1R15A Phe592 and Arg595; their catalytic efficiency (*k*_cat_/*K*_M_) was nearly 50-fold less than the wildtype, approaching that of the apo-enzyme (lacking G-actin, Fig. 2B). The more C-terminal contact with G-actin, mediated by Arg612 proved less important.

**Figure 2.**
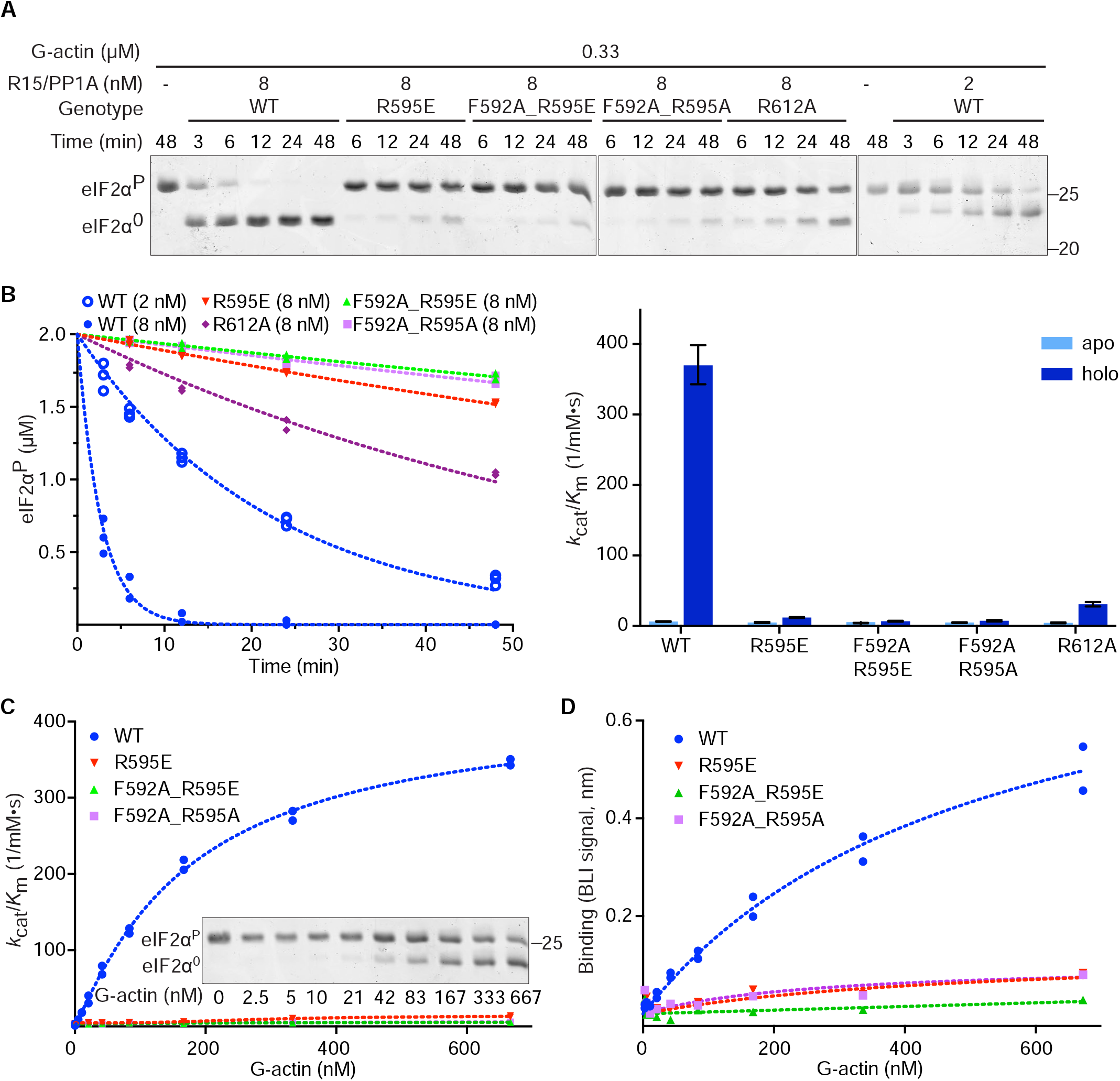
G-actin-facing PPP1R15A mutations interfere with eIF2α^P^ dephosphorylation in vitro. (A) Coomassie stained PhosTag gels resolving phosphorylated from non-phosphorylated eIF2α following eIF2α^P^ dephosphorylation in vitro with wildtype (WT) or mutant PPP1R15A holophosphatases. The concentration of binary PPP1R15A/PP1A and G-actin is indicated as is the genotype of the PPP1R15A component. Shown is a representative example of experiments reproduced at least three times. (B) **Left panel**: Time dependent progression of dephosphorylation reactions described above (showing all replicates). The dotted line is fit to a first order decay. **Right lower panel**: Mean ± 95% confidence intervals of the *k*_cat_/*K*_M_ of the indicated PPP1R15A/PP1A pairs in reactions lacking (apo) and containing (holo) G-actin. (C) Plot of the *k*_cat_/*K*_M_ of holophosphatases comprised of indicated wildtype and mutant PPP1R15A paired with PP1A as a function of G-actin concentration. A PhosTag gel (as in ‘A’) of a representative experiment with the WT enzyme is shown in the inset. (D) Plot of the association-phase plateau biolayer interferometry (BLI) signal arising from a probe comprised of immobilised wildtype or mutant PPP1R15A derivatives interacting with the indicated concentration of G-actin in solution.

The mutants’ lack of response to G-actin was confirmed in titration: Whilst binary complexes of PP1A and wildtype PPP1R15A responded to G-actin with a vigorous increase in activity, PPP1R15A^R595E^ alone, and even more so in conjunction with a PPP1R15A^F592A^ mutation, markedly attenuated the binary complex’s responsiveness to G-actin (Fig. 2C). The attenuated response of the mutant binary complexes in the enzymatic assay correlated with defective G-actin binding, as assessed by biolayer interferometry (BLI) (Fig. 2D).

This new information on functionally-important contacts between PPP1R15A and G-actin motivated us to re-visit actin’s role in eIF2α^P^ dephosphorylation in cells. PPP1R15 expression-mediated eIF2α^P^ dephosphorylation attenuates the ISR (Jousse *et al*., 2003; Novoa *et al*., 2001). This feature was exploited to examine the effect of the actin-facing PPP1R15A mutations in cells. We used a cell-based assay in which ISR activity was monitored by a CHOP::GFP fluorescent reporter (Novoa *et al*., 2001). Treatment with thapsigargin, an agent that activates the ISR by triggering PERK-dependent eIF2α phosphorylation, activated the CHOP::GFP reporter, as reflected by a shift to the right in the distribution of the fluorescent signal detected by flow cytometry. Co-expression of a control plasmid, encoding mCherry alone, had no effect on the CHOP::GFP signal. Expression of wildtype mouse PPP1R15A (fused to mCherry at its C-terminus) attenuated the thapsigargin-mediated CHOP::GFP signal (Chen *et al*., 2015). Mutant mouse PPP1R15A^R588E^ and PPP1R15A^F585A^ (counterparts to human PPP1R15A^R595E^ and PPP1R15A^F592A^) had a weaker attenuating effect on the ISR marker, a defect that was most conspicuous in the mouse PPP1R15A^F585A;R588E^, double mutant (Fig. 3A & 3B). The mouse PPP1R15A^R605A^ mutation had no observable effect in this assay, consistent with the weaker effect of the counterpart human PPP1R15A^R612A^ mutation, in vitro (Fig. 2A).

**Figure 3.**
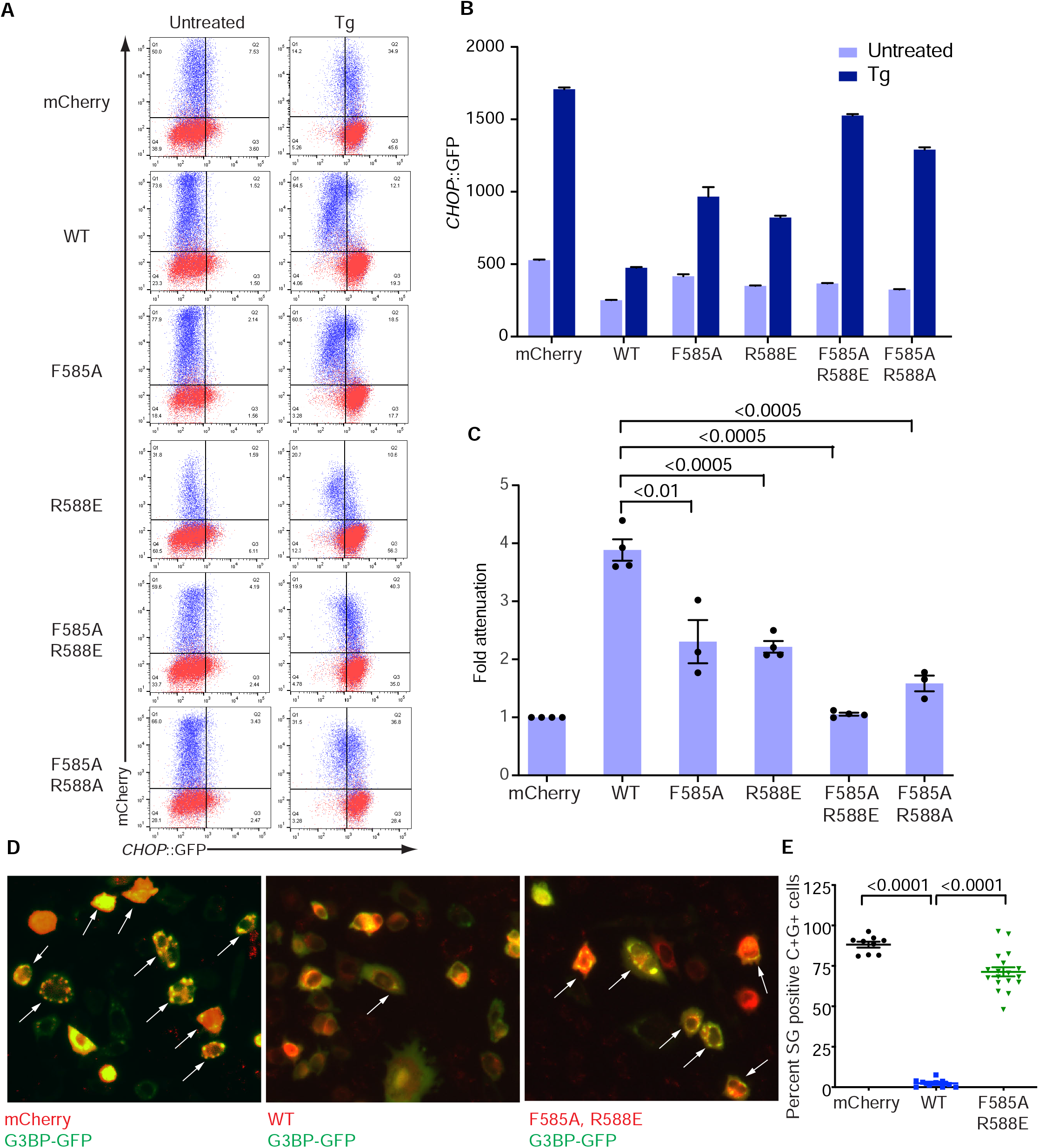
Actin facing mutations disrupt PPP1R15A action in cells. (A) Two-dimensional flow cytometry dot plots of untreated and thapsigargin (Tg)-treated CHOP::GFP transgenic CHO-K1 cells transfected with mCherry alone or full-length wildtype (WT) or mutant mouse PPP1R15A fused to mCherry at their C-termini. Note the lack of effect of mCherry on the CHOP::GFP ISR marker, the attenuation of CHOP::GFP induction by Tg in cells transfected with wildtype PPP1R15A::mCherry and defective attenuation by mutations in G-actin-facing residues in PPP1R15A. (B) The median ± SEM (n = 10^4^) of the CHOP::GFP signal in the mCherry^+^ experimental populations in ‘A’. (C) Individual data points and mean ± SEM of fold ISR attenuation by the indicated PPP1R15A::mCherry fusions from replicate experiments as in ‘A’. (P values for two-tailed T-test comparisons shown, n ≥ 3) (D) Fluorescent photomicrographs of CHO-K1 cells transfected with expression plasmids encoding the stress granule marker G3BP-GFP and mCherry alone, WT or mutant mouse PPP1R15A^F585A;R588E^::mCherry fusion proteins (as in ‘A’). Cells were treated with 0.5 mM sodium arsenite to induced stress granules (white arrows). (E) Quantitation of percent of mCherry^+^ GFP^+^ (C+G+) transfected cells with stress granules in each high-power field in sodium arsenite treated cells as in ‘D’ (P values for two-tailed T-test comparisons shown, n ≥ 8).

Stress granule formation is a convenient orthogonal marker of ISR activation (Kedersha *et al*, 1999). Consistent with this notion, their abundance in stressed cells was attenuated by enforced expression of wildtype mouse PPP1R15A, but less so by expression of PPP1R15A with G-actin facing mutations (Fig. 3D & 3E).

The effect of the actin-facing mutations on PPP1R15A action in cells is consistent with G-actin contributing to holoenzyme activity in vivo. To explore this issue further we compared β-actin that was wildtype at its PPP1R15A-contacting surface with a charge-reversal mutant β-actin^E334R^ (a residue contacting human PPP1R15A Arg595, Fig. 1C) for their ability to restore ISR inhibition when co-expressed with the severe, charge-reversed mutant mouse PPP1R15A^F585A;R588E^. To favour retention of the exogenous β-actin in its G-form, a polymerisation-deficient variant was used (β-actin^A204E;P243K^) (Joel *et al*, 2004). Neither enforced expression of otherwise wildtype nor mutant β-actin^E334R^ affected the ability of wildtype PPP1R15A to attenuate the ISR. However, β-actin^E334R^ consistently (albeit partially) reversed the defect in mouse PPP1R15A^F585A;R588E^-mediated suppression of the ISR (red arrow in Fig. 4A & 4B).

**Figure 4.**
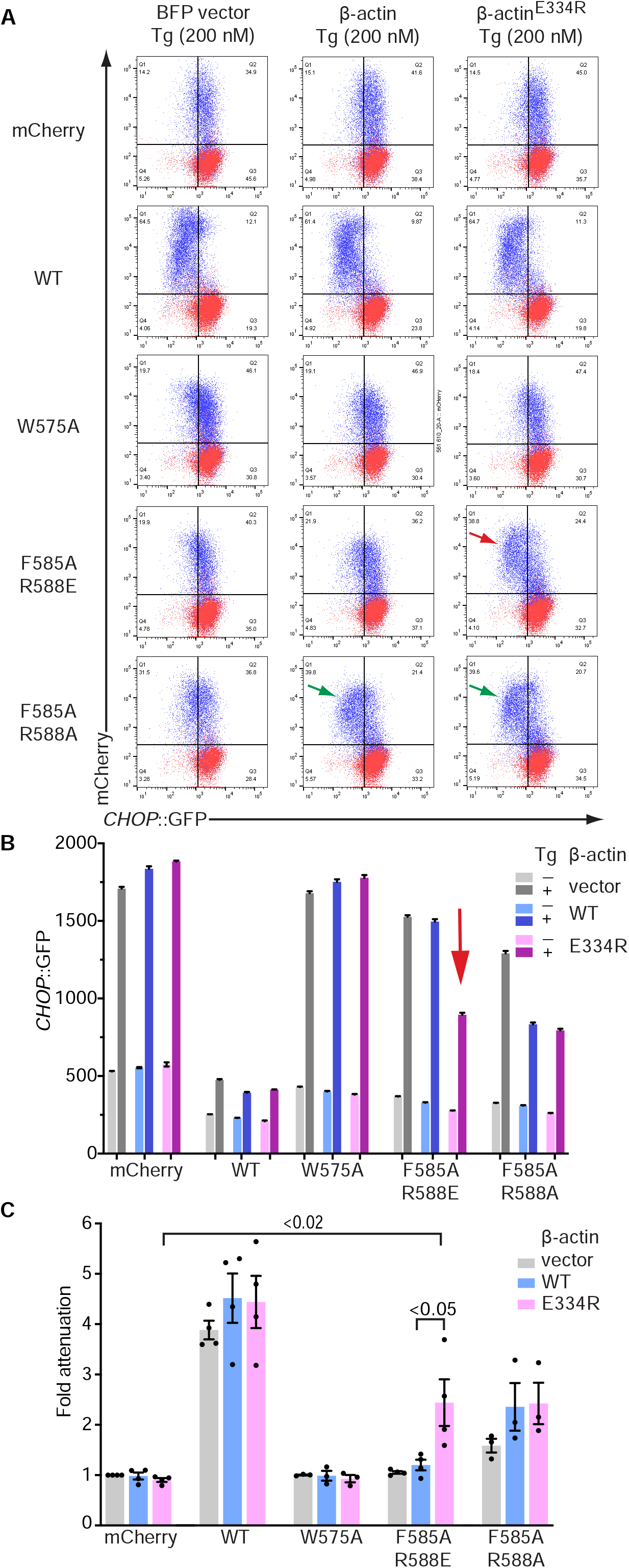
Allele-specific suppression of a PPP1R15A surface charge mutation by a reciprocal surface charge mutation in β-Actin. (A) Two-dimensional flow cytometry dot plots of thapsigargin-treated cells transfected with plasmids encoding either blue fluorescent protein (BFP) alone, BFP linked (in *trans*) to an otherwise wildtype (polymerisation-deficient A204E;P243K) β-actin or BFP linked to polymerisation-deficient β-actin^E334R^ surface-charge reversal mutant (affecting a residue that forms a salt bridge with mouse PPP1R15A^R588^). Cells were co-transfected with plasmids encoding mCherry alone or the indicated mouse PPP1R15A::mCherry fusion proteins (as in Fig. 3). Shown are the mCherry and GFP channels of the β-actin expressing (BFP^+^, blue; BFP^−^, red) populations in each data set. β-actin^E334R^ selectively restores the ability of the PPP1R15A^F585A,R588E^ mutation to attenuate the ISR (red arrow), whereas both the wildtype and β-actin^E334R^ enhance ISR attenuation by PPP1R15A^F585A, R588A^ (green arrows). (B) Display of the median ± SEM of the CHOP::GFP signal in untreated (-) and thapsigargin-treated (+) from the BFP^+^, mCherry^+^ cells in ‘A’ above (n = 10^4^). The signal in the critical sample co-expressing PPP1R15A^F585A:R588E^ and β-actin^E334R^ is denoted by a vertical red arrow. (C) Individual data points and bar diagram of the mean ± SEM ISR attenuation factor of the indicated PPP1R15A::mCherry fusions from replicate experiments as in ‘A’ (P values for two-tailed T-test comparisons shown, n ≥ 3).

Genetic rescue by the charge-substituted β-actin^E334R^ was PPP1R15A allele-specific: both wildtype and β-actin^E334R^ modestly but similarly enhanced the activity of the mouse PPP1R15A^F585A;R588A^ mutant (green arrows, Fig. 4A), whereas neither reversed the defect in the mouse PPP1R15A^W575A^ mutant (the counterpart to human PPP1R15A^W582^). These differences in response of the PPP1R15A mutants to actin are consistent with charge complementarity of the reciprocal β-actin^E334R^/PPP1R15A^R588E^ pairing and charge exclusion of the wildtype β-actin Asp^334^/PPP1R15A^R588E^ pairing accounted for the selective (albeit partial) genetic rescue in case of the former. The mouse PPP1R15A^R588A^ mutation was indifferent to the surface charge of the exogenous β-actin, accounting for the similar ability of increased concentrations of wildtype or β-actin^E334R^ to partially restore its function by mass action. The mouse PPP1R15A^W575A^ mutation, which affects PPP1R15A by a different mechanism (see below) and binds G-actin with wildtype affinity (Crespillo-Casado *et al*., 2017), was indifferent to an increase in concentration of β-actin, wildtype or mutant.

### Structure of the PP1A/PPP1R15A/G-actin-eIF2α^P^ pre-dephosphorylation complex

To better understand the catalytically-efficient tripartite holophosphatase we sought to trap a pre-dephosphorylation complex for structural analysis. To this end, we exploited observations that a PP1A^D64A^ mutation in the active site attenuates dephosphorylation and facilitates trapping of enzyme-substrate pre-dephosphorylation complexes (Wu *et al*, 2018). The substrate (the N-terminal lobe of eIF2α^P^), co-eluted with a catalytically deficient PP1A^D64A^/PPP1R15A/G-actin tripartite holophosphatase in size-exclusion chromatography and the complex gave rise to well-defined particles in Cryo-EM (Supplementary fig. 2A, supplementary fig. 2C & Table 2).

**Table 2.** Cryo-EM data collection, refinement and validation statistics.

| | PP1A<sup>D64A</sup>/PPP1R15A<sup>553-624</sup>/G-actin/DNase I/eIF2$\alpha^P$-NTD |
| --- | --- |
| **Data Collection** |  |
| Microscope | Titan Krios |
| Camera | K3 counting |
| Magnification | 130,000x |
| Voltage (kV) | 300 |
| Electron exposure (e/Å<sup>2</sup>) | 46.84 |
| Defocus range (μm) | -2.8 to -1.0 |
| Pixel size (Å) | 0.652 (counted super-resolution 0.326) |
| Symmetry imposed | C1 |
| Initial particle images | 132,495 |
| Final particle images | 60,413 |
| Map resolution (Å) | 3.96 |
| FSC threshold | 0.143 |
| Map resolution range (Å) | 3.5-6.4 |
| **Refinement** |  |
| Initial models used (PDB code) | 4MOV, 2A42 and 1KL9 |
| Model resolution (Å) | 4.1 |
| FSC threshold | 0.5 |
| **Model composition** |  |
| Non-hydrogen atoms | 9109 |
| Protein residues | 1157 |
| Waters | 0 |
| Ligands | 4 |
| **B factors (Å<sup>2</sup>)** |  |
| Protein | 47.19 |
| Ligand | 54.66 |
| **R.m.s. deviations** |  |
| Bond lengths (Å) | 0.006 |
| Bond angles (°) | 1.078 |
| **Validation** |  |
| MolProbity score | 1.81 |
| Clashscore | 10.99 |
| Poor rotamers (%) | 0 |
| **Ramachandran Plot** |  |
| Favored (%) | 96.24 |
| Disallowed (%) | 0 |
| EMDB/PDB code | EMD-12665/7NZM |

A Cryo-EM map at overall 3.96Å resolution was reconstructed and one copy of each component: PP1A^D64A^/PPP1R15A/G-actin/DNase I/eIF2α^P^, was well resolved in the map (Fig. 5A). PP1A is bound by the conserved N-terminal segment of PPP1R15A (residues 560-582 are resolved). High resolution crystal structures of PP1A/PPP1R15A (PDB 4XPN) and PP1G/PPP1R15B (PDB 4V0X) readily dock in the Cryo-EM map. The PPP1R15A chain, truncated in the crystal structure at Ala568, can be completed in the Cryo-EM map and is seen to follow the known trajectory of PPP1R1B on PP1’s surface (Supplementary fig. 2D). The crystal structure of the binary PPP1R15A/G-actin (with the bound DNase I) also docks comfortably in the Cryo-EM density and the continuity of a composite PPP1R15B>A chain is readily established in the consecutively docked crystallographic structures (Supplementary fig. 2D). eIF2α’s β-barrel engages one face of PPP1R15A’s kinked helical extension (the other face of which engages G-actin) (Fig. 5B) and the substrate loop containing pSer51 is inserted into the active site of the mutant PP1A^D64A^ (Fig. 5C & supplementary fig. 2E).

**Figure 5.**
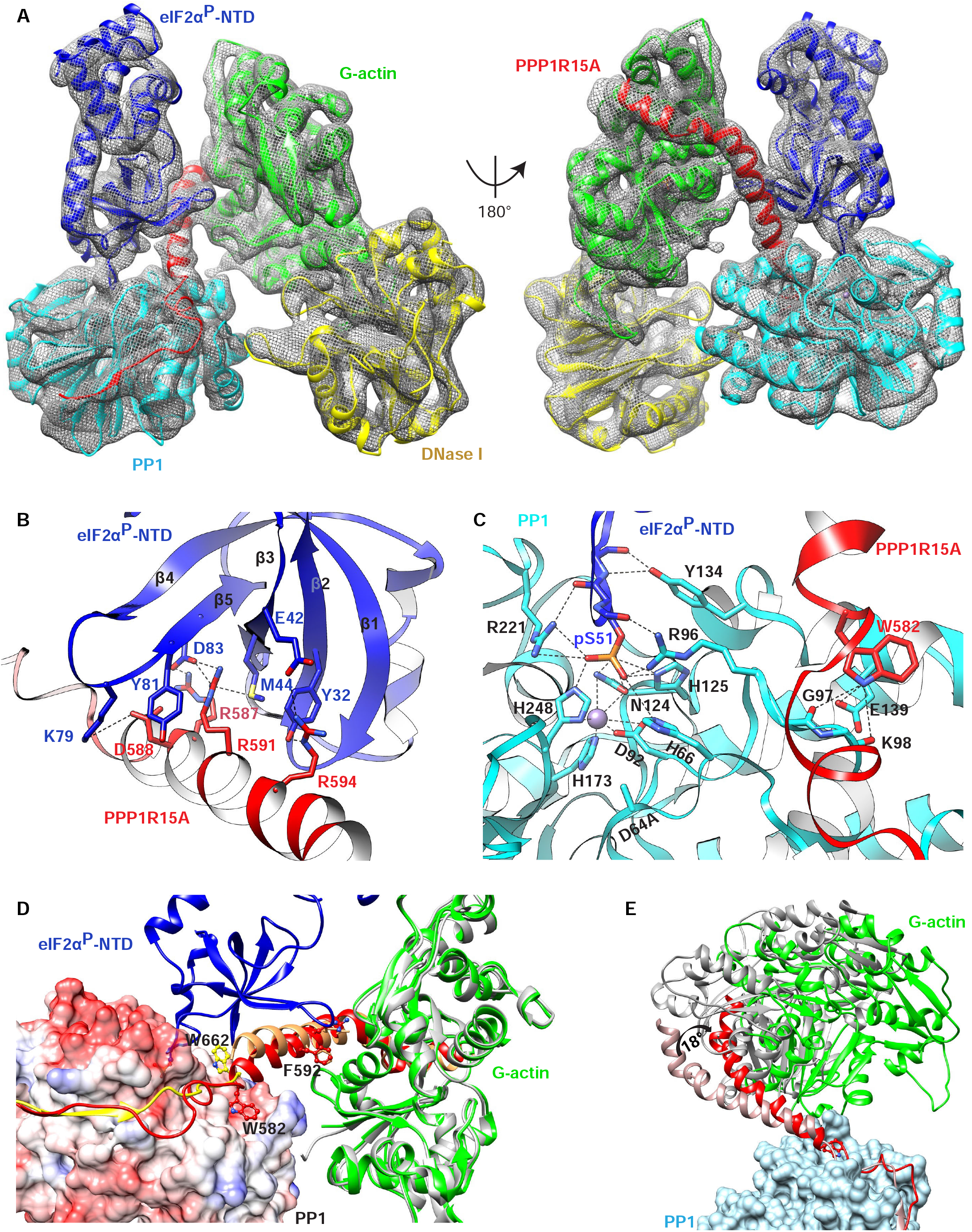
A Cryo-EM structure of the eIF2α^P^ pre-dephosphorylation complex. (A) Overview of the Cryo-EM structure of the pre-dephosphorylation complex (rendered as a GSFSC map in CryoSPARC). The threshold was set to keep the enclosed volume continuous in UCSF Chimera. Constituent proteins labelled. eIF2α^P^-NTD refers to its phosphorylated N-terminal domain. (B) Close-up view of contacts between the β-barrel of eIF2α^P^ and the non-actin-facing surface of PPP1R15A from the Cryo-EM structure. Dotted lines mark hydrogen bonding interactions. (The density map for PPP1R15A is shown in supplementary fig. 2D. (C) Close-up view of the eIF2α^P^ substrate loop (blue) in the PP1A active site (cyan). Substrate-contacting active-site residues are indicated, as eIF2α^P^ residue pS51 and the mutated PP1A^D64A^. The grey ball denotes the M2 metal ion. The density map for the eIF2α^P^ substrate loop (residues 48-53) is shown in Fig.EV2E. (D) Surface view of PP1A (coloured by charge for orientation) with PPP1R15A (in red), G-actin and eIF2α^P^ in ribbon diagram (all from the Cryo-EM structure). PPP1R15B from the binary PP1G/PPP1R15B complex (PDB 4V0X, aligned to the Cryo-EM structure by PP1G) in yellow and PPP1R15A from the crystal structure of its binary complex with G-actin/DNase I (here, aligned to the Cryo-EM structure by G-actin) in tan. Note the different disposition of PPP1R15B Trp662 (from the binary PP1G/PPP1R15B complex) and its counterpart, PPP1R15A Trp582 (from the Cryo-EM structure of the pre-dephosphorylation complex). The ‘Trp up’ disposition (exemplified by PPP1R15B Trp 662, yellow) favours the PPP1R15 helix (C-terminal to it) as found in the binary PPP1R15A/G-actin/DNase I complex (shown in tan), where it would clash with eIF2α^P^. (E) Superposition of the Cryo-EM pre-dephosphorylation complex (coloured as in ‘A’) and the crystal structure of the PP1G/PPP1R15B/G-actin complex (PDB 4V0U, in ivory and grey) aligned by PP1c. Note the ∼18° rotation of the G-actin bound PPP1R15B helical assembly, that would lead it to clash with eIF2α^P^. Supplementary movie 1 shows a back view of clashes.

Though present in the crystallised constructs, residues C-terminal to PPP1R15B Trp662 (the counterpart of PPP1R15A Trp582) were invariably disordered in binary complexes with PP1c (Chen *et al*., 2015; Choy *et al*., 2015). This suggests an important role for G-actin binding in stabilising the helical conformation of this segment. Engagement of eIF2α’s β-barrel by this helical assembly and the attendant positioning of the substrate loop in the holoenzyme’s active site provide a structural explanation for G-actin’s role as a co-factor in eIF2α^P^ dephosphorylation.

Alignment of PP1G/PPP1R15B (PDB 4V0X) to the PP1c of the pre-dephosphorylation complex reveals a conspicuous difference in the location of an invariant PPP1R15 Trp residue: in the binary complex, PPP1R15B Trp662 engages a pocket on the surface of PP1 (also bound by Phactr1 Trp542 in the PP1/Phactr1 complex, PDB 6ZEE). In the pre-dephosphorylation complex the corresponding residue, PPP1R15A Trp582, has flipped into a different position on PP1c’s surface (Fig. 5D). Enforcing the ‘Trp up’ position, observed in the binary PP1G/PPP1R15B complex on PPP1R15A in the holoenzyme, would swing the helical assembly upwards to clash with the substrate (Fig. 5D). The ‘Trp down’ position, observed in the pre-dephosphorylation complex, is stabilised locally by hydrogen bonds between the NE atom of Trp582, PP1A Asp139 side chain and the carbonyl oxygens of Gly97 and Lys98 bringing the helical assembly into a conformation compatible with substrate recruitment and catalysis. These findings provide a structural explanation for observations that though it makes no measurable contribution to PP1 or G-actin binding to PPP1R15 (Crespillo-Casado *et al*., 2017), this invariant Trp residue has an important role in substrate dephosphorylation (Chen *et al*., 2015) and below.

Plasticity of the holoenzyme is also supported by comparing the crystal structure of the PP1G/PPP1R15B/G-actin holophosphatase (PDB 4V0U) with the Cryo-EM structure of the pre-dephosphorylation complex. When aligned by their PP1s, the G-actin-stabilised helical assembly of the crystallised holoenzyme is tilted ∼18° towards the substrate (as observed in the pre-dephosphorylation complex), clashing with it (Fig. 5E & Supplementary movie 1). The pivot of the tilt corresponds to the region of PPP1R15B Trp662, though the resolution of the crystal structure is too low to assign a specific ‘up’ or ‘down’ position to its sidechain.

These observations show the potential for the holoenzyme to assume both catalytically-competent and incompetent conformations.

eIF2α^P^ binding in the competent conformation positions the pSer51 substrate loop in the PP1^D64A^ mutant active site. Density consistent with a phosphate is attached to Ser51, marking this as a pre-dephosphorylation complex. The Ser51 phosphoryl group, which occupies the position of a phosphate ion commonly observed in other PP1c structures, interacts with surrounding PP1A residues (His66, Arg96, Asn124, His125, Arg221, His248) (Fig. 5C). As expected, the M1 metal that is coordinated by PP1 Asp64 (and aligns the attacking water molecule in the wildtype enzyme (Goldberg *et al*, 1995; Wu *et al*., 2018)) is missing in this PP1A^D64A^ mutant complex. Instead, the side chain of PP1A His248 moves from its more distal position, found in bi-metallic PP1s, to contact a phosphate oxygen of eIF2α^P^ here.

There are no known structures of isolated eIF2α^P^. The substrate loop is disordered in the pre-phosphorylation complex with PKR (PDB 2A19). In complexes of eIF2(α^P^) with its nucleotide exchange factor eIF2B, pSer51 faces the interior of the loop, a conformation that may be stabilised by interactions with surrounding eIF2B residues (PDB 6O9Z, 6I3M, 6JLZ, 7D43). Ser51 is also inward facing in NMR solution structures of isolated non-phosphorylated eIF2α (PDB 1Q8K), suggesting this to be the favoured conformation of the substrate loop. Contacts observed here between loop backbone and PP1 residues Tyr134, Arg221 and Arg96 (Fig. 5C & supplementary fig. 2E) may play a role in stabilising its extended, catalytically-competent conformation, with pSer51 facing outwards, but in this, there seems no direct role for PPP1R15.

PPP1R15’s role in promoting catalysis is played out by enabling distant enzyme-substrate interactions. These are contingent on G-actin-mediated stabilisation of PPP1R15A’s helical assembly. Binding of G-actin to one face of the helix presents its other face to the conserved β-barrel of the substrate, enabling several interactions: PPP1R15A Arg591 is buried between eIF2α Met44 and Tyr81 and forms a salt bridge with Asp83. PPP1R15A Arg594 forms hydrogen bonds with eIF2α Tyr32 and a salt bridge with Asp42. PPP1R15A Arg587 forms a salt bridge with eIF2α Asp83 and eIF2α Lys79 is within range of salt bridges with PPP1R15A Asp588 and G-actin Asp25 (Fig. 5B). These substrate interactions are limited to the N-terminal half of the kinked helical assembly of PPP1R15A, which may explain the greater impact of mutations affecting N-terminal contacts with G-actin (PPP1R15A^F592A^ and PPP1R15A^R595E/A^) compared with perturbations affecting C-terminal contacts (PPP1R15A^R612A^ or cytochalasin D) (Fig. 2 & 3 and reference (Chen *et al*., 2015)).

### Functional validation of substrate recognition by the PP1A/PPP1R15A/G-actin holophosphatase

To examine the functional importance of enzyme/substrate contacts noted above, we focused on three mutations: PPP1R15A^R591A^ and PPP1R15A^R594A^, predicted on structural grounds to disrupt key contacts with eIF2α’s β-barrel, and PPP1R15A^W582A^, predicted to destabilise the active conformation of the holophosphatase. Mutation of any one of these three residues to alanine markedly weakened the holophosphatase in vitro, with PPP1R15A^R591A^ and PPP1R15A^W582A^ having the strongest effect (Fig. 6A-6C). In cells, too, the corresponding mutations, mouse PPP1R15A^R584A^ and PPP1R15^W575A^, attenuated the ability of the expressed mouse PPP1R15A to block the ISR, or interfere with stress granule formation (Fig. 6D-6G).

**Figure 6.**
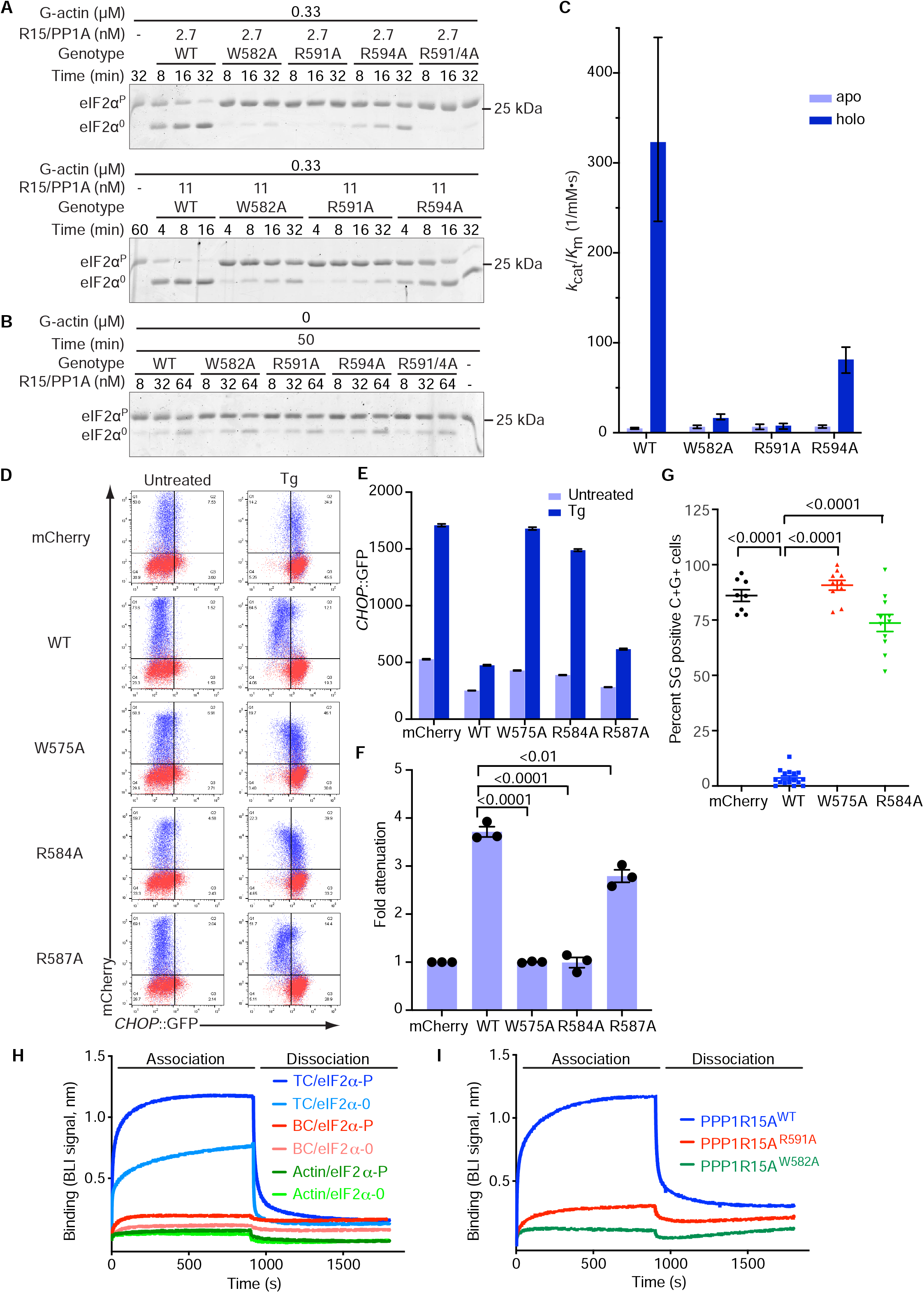
Substrate-facing PPP1R15A mutations interfere with eIF2α^P^ dephosphorylation. (A) Coomassie stained SDS-PAGE PhosTag gels of eIF2α following dephosphorylation in vitro with PPP1R15A holophosphatases. The concentration of the binary PPP1R15A/PP1A and presence of G-actin (0.33 µM) is indicated as is the genotype of the PPP1R15A component. Shown is a representative of experiments reproduced at least three times. (B) As in ‘A’ but in absence of G-actin. (C) Display of the mean ± 95% confidence intervals of the *k*_cat_/*K*_M_ of the indicated PPP1R15A/PP1A pairs in reactions lacking (apo) and containing G-actin. (D) Two-dimensional flow cytometry plots of untreated (UT) and thapsigargin (TG)-treated CHOP::GFP transgenic CHO-K1 cells transfected with mCherry alone or full-length (wildtype or mutant) mouse PPP1R15A fused to mCherry at its C-termini (as in Fig. 3A). (E) Display of the median ± SEM of the CHOP::GFP signal in untreated and thapsigargin-treated cells from ‘D’. (F) Individual data points and bar diagram of the mean ± SD CHOP::GFP attenuation factor of the indicated PPP1R15A::mCherry fusions from replicate experiments as in ‘D’ (P values for two-tailed T-test comparisons shown, N ≥ 3). (G) Quantitation of stress granules in sodium arsenite treated cells expressing the proteins as in ‘D’ (P values for two-tailed T-test comparisons shown, N ≥ 8) (H) Time-dependent change in the biolayer interferometry (BLI) signal arising from phosphorylated, eIF2α^P^, and non-phosphorylated, eIF2α^0^, immobilised on a BLI probe and introduced in a 4 µM solution of a ternary complexes (TC, PP1A^D64A^/PPP1R15A/G-actin), binary complexes of the same, lacking G-actin (BC) or G-actin alone. 900 seconds into the association phase the probe was shifted to a solution lacking protein, to record the dissociation phase. (I) BLI signal arising from immobilised eIF2α^P^ and ternary complexes of PP1A^D64A^, G-actin and wildtype or the indicated mutant PPP1R15A (as in ‘H’). Shown are representative traces of experiments reproduced at least 3 times.

The formation of a pre-dephosphorylation complex gave rise to a robust binding signal in BLI, with biotinylated eIF2α^P^ (immobilised on the probe) interacting with a mutant PP1A^D64A^/PPP1R15A/G-actin holophosphatase (in solution) (Fig. 6H). Complex formation depended on G-actin, consistent with the latter’s role in stabilising the substrate binding conformation of the holophosphatase. When used as a probe, biotinylated non-phosphorylated eIF2α^0^ also formed a complex with the holophosphatase, but it was less stable. Importantly, the PPP1R15A^W582A^ and PPP1R15A^R591A^ mutations, that interfered with the enzyme’s catalytic efficiency also interfered with substrate binding in BLI (Fig. 6I).

Contacts present in the pre-dephosphorylation complex involve eIF2α residues previously-observed to interact with the kinase PKR in a pre-phosphorylation enzyme/substrate complex (Dar *et al*., 2005). Notably, the sidechain of PKR Phe489 is inserted into the cleft formed by eIF2α Met44 and Tyr81 (a counterpart to PPP1R15A Arg591, whose mutation blocked dephosphorylation, Fig. 6A). eIF2α Lys79 forms a salt bridge with PKR Glu490 in the pre-phosphorylation complex and to be within range of salt bridge formation with PPP1R15A Asp588 and actin Asp25, in the pre-dephosphorylation complex, whereas eIF2α Asp83 hydrogen bonds with PPP1R15A Arg591 in the pre-dephosphorylation complex and with the backbone nitrogen of PKR Ala488 in the latter’s substrate-binding helix G (Fig. 7A and reference (Dar *et al*., 2005)).

**Figure 7.**
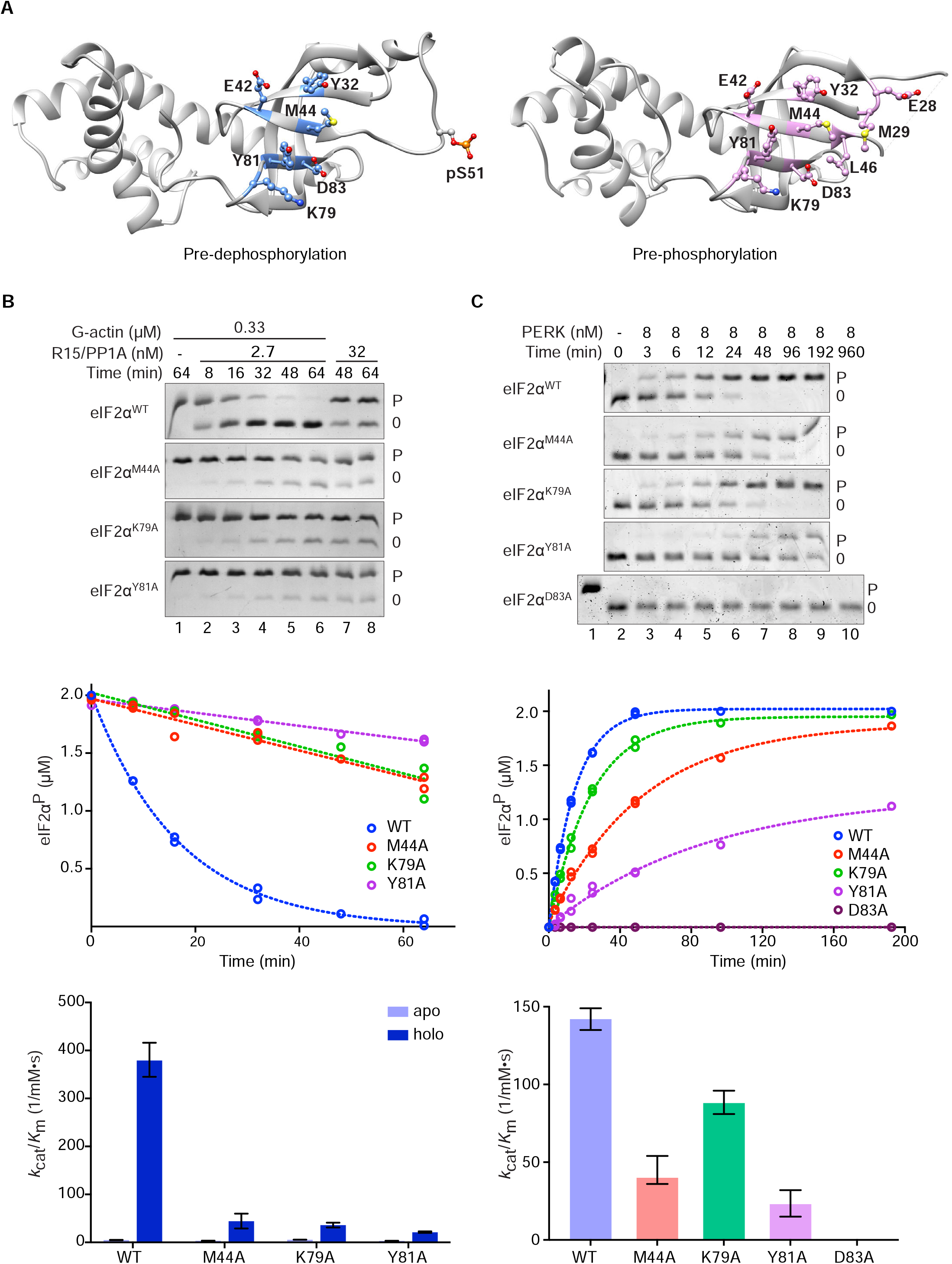
Enzyme-facing substrate mutations interfere with eIF2α^P^ dephosphorylation. (A) Ribbon diagram of eIF2α from the pre-dephosphorylation complex (from here, **left**) and a pre-phosphorylation complex with PKR (PDB 2A19, **right**). Residues contacting the respective enzymes are highlighted. pSer51, shown on the left, was unresolved in PDB 2A19. (B) **Upper panel**: Coomassie stained PhosTag gels of wildtype and the indicated eIF2α^P^ mutants (at 2 µM), following dephosphorylation in vitro with PPP1R15A holophosphatases. **Middle panel**: Time dependent progression of the dephosphorylation reactions above. The dotted line is fit to a first order decay (of replicates). **Lower panel**: Mean ± 95% confidence intervals of the *k*_cat_/*K*_M_ of the indicated enzyme/substrate pairing in reactions lacking (apo) and containing (holo) G-actin. Shown are representative example of experiments reproduced at least three times. (C) As in ‘B’ (above) but reporting of the phosphorylation of the wildtype and the indicated eIF2α^0^ mutants (at 2 µM) by the indicated concentration of the kinase PERK.

In keeping with the functional importance of these contacts, we observed that eIF2α^M44A^ and eIF2α^Y81A^ mutations interfered with its ability to serve as a substrate of both the PPP1R15A-containing holophosphatase and the ISR inducing kinase PERK, in vitro. The eIF2α^K79A^ mutation also affected both reactions; de-phosphorylation more than phosphorylation (Fig. 7B & 7C). The effect of the eIF2a^D83A^ mutation on dephosphorylation could not be tested as it blocked all phosphorylation (Fig. 7C). Together these observations point to convergence of higher order enzyme/substrate contacts in opposing reactions affecting eIF2α.

## DISCUSSION

The structure of a PPP1R15A-based pre-dephosphorylation complex presented here rationalises G-actin’s role as an essential component of the holophosphatase in vitro. Corroborating biochemical studies and allele-specific suppression of a defective PPP1R15A by a rationally-designed compensatory mutation in β-actin, link G-actin to holophosphatase function in vivo, too. Actin-dependent contacts between the holophosphatase and a distant surface of eIF2α position pSer51 at the enzyme’s active site, paralleling distal contacts conserved in the pre-phosphorylation complex. Thus, the kinases that initiate and the phosphatases that terminate signalling in the ISR appear to have converged on a common solution for catalytic efficiency, one that relies on recognising features of their folded globular substrate, distinct from its substrate loop.

The enzyme studied here contains the conserved core, common to PPP1R15s from all phyla. Likewise, the independently folded N-terminal domain of eIF2α, which extends flexibly from the eIF2 trimer, serves as a minimal specific substrate for kinases and phosphatases (Ito *et al*, 2004). Co-factors other than G-actin may contribute to PPP1R15A-mediated dephosphorylation in cells. Interactions between the extended N-termini of PPP1R15A (or B) and the β or γ subunits of the eIF2(α^P^) trimer (missing here) may contribute further to catalysis. But for now, these are unproven. Furthermore, the biochemical properties of the minimal system studied here in vitro, are adequate to explain the kinetics of eIF2α^P^ dephosphorylation observed in cultured cells (see *enzymology* in methods section). Therefore, the enzyme described here is a valid starting point to study the dephosphorylation event that terminates signalling in the ISR.

The observation that higher-order contacts distant from both the substrate loop and enzyme active site promote dephosphorylation, presents a challenge: whilst affinity of an enzyme for its substrate favours catalysis, residual affinity for the product is often anti-catalytic. This conundrum could be settled by dephosphorylation-dependent structural changes to eIF2α that favour product dissociation. Allosteric changes have been noted in eIF2α upon Ser51 phosphorylation/dephosphorylation (Kashiwagi *et al*, 2019; Kenner *et al*, 2019), but they do not extend to the region involved in the higher order contacts identified here. Furthermore, given the similarity in contacts with the kinases that phosphorylate eIF2α^0^ and the PPP1R15-holophosphatase that dephosphorylates eIF2α^P^, a coherent allosteric change that promotes product release in both antagonistic reactions seems unlikely.

In an alternative scenario, cooperation between local contacts at the active site and higher order contacts is important in formation of the pre-reaction complex. Enhanced binding of the holophosphatase to immobilised eIF2α^P^, compared with its binding to non-phosphorylated eIF2α (in the BLI assay), speaks to this point. The PP1A^D64A^ mutation, used to stall the reaction, destabilises the M1 metal ion of the active site, which can no longer coordinate an eIF2α pSer51 oxygen (Goldberg *et al*., 1995). Due to loss of these contacts, the difference in affinity of the substrate (eIF2α^P^) and product (eIF2α^0^) for the wildtype holoenzyme may be greater than that reported by the BLI experiment; possible great enough to dominate the kinetics of product release. The higher order contacts are clearly important, as both binding in the BLI assay and the 50-fold spread in catalytic efficiency between the apo and holophosphatase are sensitive to their disruption. However, it stands to reason that successful product dissociation requires that the affinity imparted by these distal contacts be tuned to differences in binding kinetics of the active site for the substrate versus product. The higher off rates of the product observed in BLI likely play into this. Similar considerations apply to eIF2α phosphorylation, in which higher order contacts likely cooperate with contacts at the active site to stabilise a pre-dephosphorylation complex (Dar *et al*., 2005; Marciniak *et al*, 2006). We speculate that the relatively modest surface of eIF2α buried in both complexes (∼1200 Å^2^) reflects a constraint imposed on substrate and product affinity by catalytic efficiency.

PPP1R15’s positive role in eIF2α^P^ dephosphorylation appears limited to scaffolding the enzyme to favour the aforementioned higher order contacts - PPP1R15A makes no contacts with the eIF2α^P^ substrate loop. Nonetheless, local sculpting of PP1c’s surface by regulatory subunits is likely to affect eIF2α^P^ dephosphorylation. For example, insertion of the eIF2α^P^ substrate loop in PP1c’s active site is compatible with neither Phacter1 nor spinophilin binding (Fedoryshchak *et al*., 2020; Ragusa *et al*., 2010). Thus, the process described here for selecting a globular domain of a phosphoprotein as substrate liklely operates alongside the well-established mechanism for locally biasing access to the PP1c active site by regulatory subunit engaging its surface (Peti *et al*., 2013).

In eIF2α phosphorylation, formation of higher-order contacts with the substrate is dependent on allosteric coupling of back-to-back dimerisation of the kinase to alignment of its eIF2α-binding helix G (Dey *et al*, 2005). Precise alignment of components is also required for higher-order contacts in the tripartite holophosphatase. This fine tuning seems to have arisen solely by refinement of the PPP1R15 component, as features that enable it to function together with PP1c and G-actin are present in simpler eukaryotes that have no counterpart to PPP1R15. For example, mammalian actin Asp334 that contacts PPP1R15A Arg595 is conserved in yeast. Similarly conserved is the actin hydrophobic pocket that accommodates PPP1R15A Phe592. Indeed, PPP1R15A is functional as an eIF2α^P^ phosphatase in yeast and its activity depends on integrity of residues involved in what we now understand to be G-actin contacts (Rojas *et al*, 2015). It thus appears that in metazoans the holophosphatase was cobbled together from one rapidly evolving component (PPP1R15) and two off-the-shelf pre-existing blocks (PP1 and G-actin).

Actin is a highly dynamic protein and its partitioning between filamentous, F-actin and monomeric, G-actin, is responsive to upstream signals. G-actin is a limiting ligand in some physiological reactions, subordinating them to the F/G-actin ratio (Mouilleron *et al*, 2008). Actin dynamics may have been co-opted to regulate the ISR in some circumstances, but these have yet to be identified. Alternatively, G-actin incorporation into the holophosphatase may have arisen simply by its availably as a convenient building block. The genetic evidence provided here for G-actin’s role as a co-factor in eIF2α^P^ dephosphorylation in cells renders these questions pertinent.

Potent inhibitors of PP1c exist, but they non-selectively block dephosphorylation. Mechanism-based pharmacological targeting of substrate-specific holophosphatases is a long-sought goal, which, despite claims to contrary (Carrara *et al*, 2017), may have yet to be realised (Crespillo-Casado *et al*., 2017; Crespillo-Casado *et al*., 2018). Dependence of the eIF2α^P^ dephosphorylation reaction on the precise alignment of the three components of the holophosphatase hints at a path towards this goal. As noted above, the catalytic and substrate binding lobes of the holophosphatase pivot about a hinge located between the PP1-facing and G-actin facing portions of PPP1R15. In the catalytically-favoured conformation the conserved PPP1R15A Trp582 may transition from its conventional pocket on the surface of PP1A (observed in the binary PP1A/PPP1R15B and PP1A/Phacter1 complexes (Chen *et al*., 2015; Fedoryshchak *et al*., 2020)) to a novel pocket observed here. Ligands that bind the holophosphatase and stabilise the inactive conformation of Trp582 are predicted to selectively inhibit eIF2α^P^ dephosphorylation by interfering with the higher-order contacts necessary for terminating signalling in the ISR. Time will tell if such insights, derived from these snapshots of the eIF2α^P^ phosphatase in action, advance targeting of dephosphorylation reactions to beneficial ends.

## MATERIALS AND METHODS

### Protein expression and purification

N-terminal His6-Smt3 human PPP1R15A^582-621^ (UK2514) used for co-crystallisation with G-actin and DNase I was expressed in *E. coli* BL21 T7 Express *lysY/I^q^* cells (New England BioLabs, cat. # C3013). The bacterial cultures were grown at 37 °C to an optical density (OD600) of 0.8 in 2 × TY medium supplemented with 100 μg/ml ampicillin, and expression of recombinant protein was induced at 22 °C for 16 h by the addition of 0.5 mM isopropylthio β-D-1-galactopyranoside (IPTG). After harvesting by centrifugation, the pellets were suspended in the HisTrap column-binding buffer (20 mM Tris-HCl pH 7.4, 0.5 M NaCl, 20 mM imidazole) containing protease inhibitors and 0.1 mg/ml DNase I and 20ug/ml RNase A. The cells were crushed by a cell disruptor (Constant systems) at 30 kPSI and the obtained lysates were cleared by centrifugation for 1 h at 45,000× g. The supernatant was applied to a pre-equilibrated 5 ml HisTrap column (GE Healthcare) using a peristaltic pump. After washing with about 100 ml of the binding buffer, the bound fusion protein was eluted with 20– 200 mM imidazole gradient using a FPLC purifier system (ÄKTA; GE Healthcare). Peak fractions were pooled and digested with SENP2 protease (at final 0.01 mg/ml; produced in-house) to cleave off the His6-Smt3 tag overnight at 4 °C. After complete digestion, the remaining full-length fusion protein and His6-Smt3 tag were removed by binding back to a 5ml HisTrap column. Following buffer exchange to lower salt buffer (10 mM HEPES–KOH pH 7.4, 50 mM NaCl), the intact protein was further purified by cation exchange chromatography using a HiTrap SP HP column (GE Healthcare) and eluted by the 50– 500 mM NaCl gradient in 10 mM HEPES–KOH pH 7.4. Samples of fractions from each purification step were analyzed by SDS-PAGE and Coomassie staining. Target protein contained peak were collected and were further purified using a HiLoad 16/600 Superdex 75 prep grade gel filtration column equilibrated with 10 mM Tris-HCl pH 7.4 and 0.15 M NaCl. The elution peak fractions were pooled and ready for forming complex with G-actin/DNase I. Extra samples were snap-frozen in liquid nitrogen and stored at −80 °C.

H6-Smt3-tagged human PPP1R15A^553-621^ with a C-terminal fused maltose binding protein (MBP) tag and rabbit PP1A^D64A^ (7-300, UK2720) were co-expressed using the bicistronic construct with a Shine-Dalgarno (SD) sequence inserted between the 3’ of PPP1R15A and the 5’ of PP1A coding sequences in *E. coli* BL21 T7 Express *lysY/I^q^* cells, induced as described above and purified by sequential NiNTA affinity chromatrography, His6-Smt3 tag cleavage, reverse NiNTA affinity chromatrography (to remove the H6-SUMO3) and a HiLoad 16/600 Superdex 200 size-exclusion chromatography, as described above. 1 mM MnCl_2_ was included in all of the steps from IPTG induction. Sequential nickel chelating, His6-Smt3 tag cleavage and reverse nickel chelating were applied for making PP1A/PPP1R15A (UK2699), PP1A/PPP1R15A^R595E^ (UK2750), PP1A/PPP1R15A^F592A, R595A^ (UK2751), PP1A/PPP1R15A^R612A^ (UK2752) and PP1A/PPP1R15A^F592A, R595E^ (UK2755) for the dephosphorylation assay (Fig. 2A-2C).

N-terminal H6-Smt3 human eIF2α^2-187^ (eIF2α-NTD, UK2731) was purified by sequential nickel chelating, His6-Smt3 tag cleavage, reverse nickel chelating, as described above. The crude eIF2α-NTD sample was phosphorylated by incubating with GST-PERK (1:100 molar ratio) for 2h at 37 °C in the buffer of 100 mM NaCl, 25 mM Tris-HCl pH 7.4, 5 mM MgCl_2_, 2.5 mM ATP and 1 mM TCEP. After phosphorylation, the sample was further purified by a HiLoad 16/600 Superdex 75 size-exclusion chromatography equilibrated with 10 mM Tris-HCl pH 7.4, 0.15 M NaCl and 1 mM TCEP. Fractions were pooled and ready to form complexes with holophosphatase components.

Mutant derivatives of H6-Smt3 human eIF2α^2-187^ (UK2849-52) were expressed as described above, but to compensate for their defective ability to serve as substrates of phosphorylation, the reaction was extended overnight at room temperature: Substrate, at ∼50 µM, was combined with purified GST-PERK at 10 nM in: 100 mM NaCl, 25 mM Tris-HCl pH 7.4, 5 mM MgCl_2_, 2.5 mM ATP and 1 mM TCEP and incubated 16 hours at room temperature. Completion of the reaction was verified by PhosTag SDS-PAGE and residual enzyme was removed by flowing the sample through a glutathione-sepharose resin.

Biotinylated, AviTagged human eIF2α^2-187^ (UK2733, Fig. 6F) (Zyryanova *et al*, 2021) was purified and phosphorated as described above, and biotinylated AviTagged PPP1R15A^583-621^ (wildtype and mutants, UK2773,6,7,9, Fig. 2D) from cultures supplemented with Biotin, 100 µM

Actin was purified from rabbit muscle as described (Pardee & Spudich, 1982), dialyzed against G buffer (5 mM Tris-HCl pH 8, 0.2 mM ATP, 0.5 mM DTT, 0.2 mM CaCl_2_) for 3 days. The G-actin in the binary structure was further incubated with a ten-molar excess of Latrunculin B (#ab14491, abcam). Partially purified (DP grade) bovine pancreatic DNase I powder was purchased from Worthington Biochemical Corporation (#LS002139, Lakewood NJ). The DNase 1 powder was dissolved in 10 mM Tris 7.4, 0.05 M NaCl, 1 mM CaCl_2_ and protease inhibitors before purified by a HiLoad 16/600 Superdex 75 prep grade gel filtration column equilibrated with 10 mM Tris-HCl pH 7.4, 0.15 M NaCl and 1mM CaCl_2_. Peak fractions were pooled with addition of protease inhibitors. 1:1.5 molar ratio of G-actin and DNase1 were mixed before loaded to a HiLoad 16/600 Superdex 75 prep grade gel filtration column equilibrated with 10 mM Tris-HCl pH 7.4, 0.15 M NaCl and 1 mM CaCl_2,_ 0.2 mM ATP and 0.2 mM TCEP. The G-actin/DNase I complex peak fractions were collected and followed by forming complex with either human PPP1R15A^582-621^ or PP1A^D64A^/PPP1R15A^553-624^_MBP/eIF2α-NTD^P^. The former complex was further purified by a HiLoad 16/600 Superdex 75 prep grade gel filtration column equilibrated with 10 mM Tris-HCl pH 7.4, 0.15 M NaCl and 1 mM CaCl_2,_ 0.2 mM ATP and 0.2 mM TCEP. The latter complex was further purified by consecutive Superdex Increase 75 10/300 GL and Superdex Increase 200 10/300 GL gel filtration columns equilibrated with 10 mM Tris-HCl pH 7.4, 0.15 M NaCl, 1 mM CaCl_2,_ 0.2 mM ATP, 0.2 mM TCEP and 1 mM MnCl_2_. The G-actin/DNase 1/PPP1R15A^582-621^ complex peak fractions were concentrated to 10mg/ml for crystallisation; the PP1A^D64A^/PPP1R15A^553-624^_MBP/eIF2α^P^-NTD/G-actin/DNase 1 complex was concentrated to 5mg/ml for making Cryo-EM grids.

### Structural analysis

#### Crystallography

100nl of PPP1R15A^582-621^/G-actin/DNase I complex at 10mg/ml and 100nl well solution was added to each drop of 96-well sitting-drop crystallisations trays for broad screening and the crystallisation took place at 20°C. The initial thin plate crystals grew at 8% PEG4000 and 0.1 M Na acetate pH 4.6. The best dataset was collected from a crystal grew in 10% PEG4000 and 0.1 M Na acetate pH 5 microseeded with crashed initial crystals. Diffraction data were collected at beamline i03 in the Diamond Synchrotron Light Source (DLS) and processed by the XIA2 pipeline implementing DIALS (Winter *et al*, 2018) for indexing and integration, Pointless (Evans, 2011) for space group determination, and Aimless (Evans & Murshudov, 2013) for scaling and merging. The structure was solved by molecular replacement using Phaser (McCoy *et al*, 2007) and 2 copies of G-actin/DNase I (PDB 2A42) were found in an asymmetric unit. PPP1R15A^582-621^ was manually built to the difference map in COOT (Emsley *et al*, 2010). Further refinement was performed iteratively using COOT, phenix.refine (Afonine *et al*, 2018) and refmac5 (Murshudov *et al*, 2011). MolProbity (Chen *et al*, 2010) was consulted for structure validation (Supplementary Table 1). Molecular graphics were generated using UCSF Chimera (Pettersen *et al*, 2004) and PyMOL (The PyMOL Molecular Graphics System, Version 1.3 Schrödinger, LLC.).

#### Cryo-EM

UltrAuFoil 0.6/1 300 mesh grids (Quantifoil) were glow discharged in residual air at 25 mA for 3 min using a Pelco EasiGLOW^TM^. 5 μl of PP1A^D64A^/PPP1R15A^553-624^_MBP/eIF2α-NTD^P^/G-actin/DNase I complex at 5 mg/ml was mixed with 0.2 μl of 5.6 mM Triton X-100 immediately before plunging. The additional ∼0.22 mM Triton X-100 has improved particle orientation distribution significantly. 3.5 μl sample was deposited to grids and fast frozen in liquid ethane cooled by liquid nitrogen using Vitrobot Mark IV (Thermofisher). Grids were blotted for 3.5 s with a force at −7 in 100% humidity chamber at 4 °C. Movie stacks were collected on a Titan Krios operated at 300 keV equipped with a K3 camera (Gatan) at the super resolution counting mode. Images were recorded at 130,000x magnification corresponding to 0.652 Å per pixel (counted super resolution 0.326 Å per pixel) using a 20 eV energy filter. Image stacks have 49 frames for an accumulated does of ∼50 e Å^−2^ in a total exposure time of 1.3 s. The defocus range was from −2.8 μm to −1 μm. 4025 micrographs were collected from a single grid.

WARP (Tegunov & Cramer, 2019) was used for motion correction and CTF estimation, during which the super resolution data were binned to give pixel size of 0.652 Å. Particles were automatically picked using the BoxNet algorithm of WARP. 132,495 particles stacks generated by WARP were imported to CryoSPARC (Punjani *et al*, 2017). Four initial models were generated by initial 3D reconstruction from scratch, followed by a heterogeneous refinement with the four initial models and the whole particle set as input. Model 1 (29879 particles) represented G-actin/DNase I complex; model 2 and 3 were ice or unknown junks and model 4 (76280 particles) represented the full complex. Another two rounds of ab-initio 3D reconstruction and heterogeneous refinement of the model 4 particle stacks were performed to remove extra bad particles, resulting in 60413 final particles for the full complex. Further 3D classification of these 60413 particles generated two similar classes with worse resolutions at the core region, which had noticeable differences only at the intensities of the flexible MBP tag density. Non-uniform refinement (Punjani *et al*, 2020) was performed to improve the resolution, resulting a 3.96 Å map with corrected auto-tightened mask at gold standard FSC of 0.143. Defocus, global CTF and local refinement did not improve the resolution. ResLog plot (Stagg *et al*, 2014) of resolutions against particles reached plateau with the final particle set.

Docking of PP1c (PDB 4MOV), G-actin/DNase I (PDB 2A42) and eIF2α-NTD (PDB 1KL9) to the Cryo-EM map was performed in UCSF Chimera. COOT was used for further local model building. A ResolveCryoEM map (Terwilliger *et al*, 2020) was consulted to build regions with better quality, such as the C-terminal helical domain of PPP1R15A, the pSer51 loop and β-barrel of eIF2α^P^-NTD. The atomic model was refined with Phenix real-space refinement (Afonine *et al*., 2018) using the ResolveCryoEM map. MolProbity was used for structural validation (Supplementary Table 2). UCSF Chimera was used to prepare graphic figures. The angular distribution was generated by the final non-uniform refinement in CryoSPARC and converted by UCSF pyem (Asarnow et al. 2019) to plot in UCSF Chimera.

### Enzymatic activity

#### Measuring eIF2α^P^ dephosphorylation rates

Dephosphorylation reactions were conducted essentially as described (Chen *et al*., 2015). Binary complexes of wildtype or the indicated mutants of human PPP1R15A^553-621^ (fused to *E. coli* MBP at their C-termini, as a solubility tag) with associated untagged rabbit PP1A^7-300^, purified from *E. coli* as described above and maintained as concentrated stocks (≥10 µM), were quickly diluted into assay buffer (100 mM NaCl, 25 mM Tris-HCl pH 7.4), 200 µM MnCl_2_, 1 mM TCEP, 5% glycerol, 0.02% Triton X-100) to the indicated concentrations (2 – 100 nM) in a 200 µl PCR tube maintained at 20 °C. G-actin was added (to a final concentration 330 nM, unless indicated differently) and the sample was incubated at 20 °C for 5 minutes to allow ternary complex assembly. The reaction was initiated by adding phosphorylated human eIF2α^2-187^ (purified from *E. coli* and phosphorylated in vitro with GST-PERK, as indicated above) to a final concentration of 2 µM.

Samples were removed from the reaction at intervals, quenched into sample buffer (62.5 mM Tris-Glycine pH 6.8, 50 mM DTT, 2% SDS, 10% glycerol), loaded onto a 10 × 0.1 cm 15% SDS-PAGE gel with 50 µM PhosTag reagent and 100 µM MnCl_2_, resolved at 200 V for 80min, stained with Coomassie and fluorescently scanned on a Licor Odyssey using the built-in 680 nm laser.

The intensity of the phosphorylated and non-phosphorylated species in each lane were quantified using NIH image (Schindelin *et al*, 2012).

#### Measuring eIF2α phosphorylation rates

Human eIF2α^2-187^, final concentration 2 µM, and GST-PERK, final concentration 8 nM (both purified from *E. coli*) in kinase buffer (100 mM NaCl, 25 mM TrisHCl pH 7.4, 5 mM MgCl_2_, 2.5 mM ATP, 1 mM TCEP, 5% glycerol, 0.02% Triton X-100) were combined in a PCR tube at 20°C and reaction progress was monitored by SDS-PAGE PhosTag electrophoresis and Coomassie stain, as described above.

### Biolayer interferometry

BLI experiments were conducted at 30 °C on the FortéBio Octet RED96 System, at an orbital shake speed of 600 rpm, using Streptavidin (SA)-coated biosensors (Pall FortéBio) in an assay buffer of 100 mM NaCl, 25 mM Tris-HCl pH 7.4, 200 µM MnCl_2_, 1 mM TCEP, 5% glycerol, 0.02% Triton X-100.

Biotinylated ligands, either wildtype or mutant PPP1R15A^583-621^ (Fig. 2D) or phosphorylated or non-phosphorylated eIF2α^2-187^ (Fig. 6F) both expressed in and purified from *E. coli* as C-terminally-AviTag-His6-tagged proteins were loaded onto biosensor at a concentration of 150 nM to a binding signal of 1-2 nm, followed by baseline equilibration in buffer. Association reactions with analyte: G-actin (Fig. 2D) or wildtype or mutant complexes of PPP1R15A^553-624^ (fused to MBP at its C-term) with PP1A^D64A^ with or without G-actin (Fig. 6F) in the assay buffer described above were conducted with a reaction volume of 200 μL in 96-well microplates (greiner bio-one).

### ISR activity in cells

#### Cell culture & Transfection

CHOK1 wildtype (stress granule experiments) or C30 CHOP::GFP (Novoa *et al*., 2001) (Flow cytometry experiments) cells were maintained in Hams’ F12 supplemented with 10% Fetalclone II serum (Hyclone) and 1x pen-strep. For Flow cytometry, cells were seeded at a density of 1.5×10^5^ in 12 well dishes and 24 hours later were co-transfected with 50 ng mCherry, R15A-mCherry wildtype or mutant expression vector and 950 ng of BFP marked empty or Actin (WT or E334R) expression vector (1:19 ratio of R15::Actin) using Lipofectamine LTX (Life Technologies) at a ratio of 3 μl Lipofectamine LTX and 1 μl of Plus reagent per ng of DNA in following complex formation in OPT-MEM according to the manufacturer’s instructions. 12 hours post transfection the media was changed and thapsigargin added at 200 nM where indicated. The cells were washed and released from the dishes in PBS-4mM EDTA with 0.5 % BSA and 10000 BFP+ cells subject to Flow cytometry analysis as described in (Chen *et al*., 2015).

Experiments tracking formation of stress granules were adapted from (Panas *et al*, 2019) CHO-K1 cells were transfected on poly-lysine coated multiwell slides. DNA complexes composed of 100 ng each of G3BP-GFP and R15A-mCherry expression vector and 450ng of empty plasmid carrier were formed in a ratio of 3 μl/ng DNA and 1 μl of Plus reagent per ng of DNA in following complex formation in OPT-MEM according to the manufacturer’s instructions. After 15 minutes the complexes were diluted into 0.5 ml CHO medium and 125 μl aliquoted into 4 wells containing cells. Medium was changed 16 hours post transfection and 4 hours later replaced with medium containing 0.5 mM sodium arsenite. Cells were then washed and fixed in PBS-4% paraformaldehyde followed by washing and mounting for microscopy. Micrographs of the red and green channels were taken on an EVOS inverted photo-microscope using the 20x lens.

## QUANTIFICATION AND STATISTICAL ANALYSIS

### Enzyme kinetics

At concentrations well below the substrate *K*_M_ the conversion to product (i.e. substrate consumption) in an enzymatic reaction follows first order kinetics and can be described ideally a monophasic exponential decay.

When [S] << K_M_, enzymatic reactions are first order (their rate is proportional to [S]. This follows from Michaelis Menten

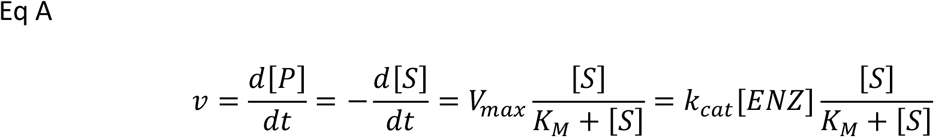

If [S] << K_M_ then this rearranges to:

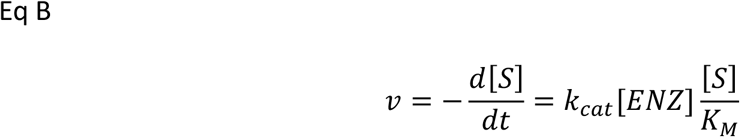

Solved for *k*_cat_/*K*_M_ this rearranges to:

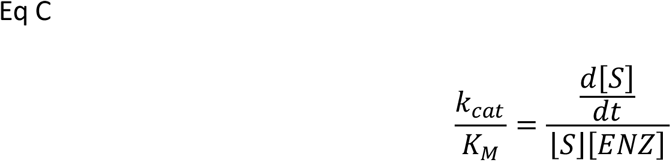

The experimental *k*_obs_ of the conversion of substrate to product (fit to a monophasic exponential decay) corresponds to the term

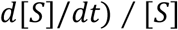

*k*_cat_/*K*_M_ can therefore be extracted from the experimentally determined *k*_obs_ and the known concentration of enzyme [*ENZ*]

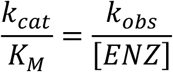

At 2 µM, eIF2α is well below the substrate *K*_M_ of both the PPP1R15 holophosphatase and

GST-PERK (reference (Chen *et al*., 2015) and data not shown). Therefore, the time dependent substrate depletion in the phosphorylation and dephosphorylation reactions was fitted to a one phase decay function on Graphpad Prism V8 using the model below:

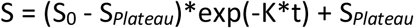

(S_0_ was constrained to the concentration of substrate at t = 0, S*_Plateau_* was set to zero as at t = ∞, S = 0)

Due to practical limitations in the time range over which one can extend the dephosphorylation assay, or accurately obtain soundings of its progression, the range over which we can measure *k*_obs_ is limited. To measure reaction progression accurately with the faster holophosphatase(s) and the much slower apo-enzymes we designed the assays to have similar *k*_obs_ by varying [ENZ]: In assays of the various holophosphatases [ENZ] = 2 nM, whereas in assays of the apo enzyme [ENZ] may be 25 times higher. Therefore *k*_cat_/*K*_M_ (which is simply *k*_obs_/[ENZ]) emerges as useful metric to compare catalytic efficiency of the holo- and apo-enzymes measured in assays that have very different [ENZ]. However, as the experiments most critical to the conclusions drawn here, comparing dephosphorylation by different holophosphatases, were conducted with the *same* enzyme concentrations [ENZ] and same initial substrate concentrations [S]_0_, and as *k*_cat_/*K*_M_ = *k*_obs_/[ENZ], the fold difference in *k*_cat_/*K*_M_ of such reactions (reported on in Fig. 2B, 6C, 7B & 7C) is identical to the fold difference of their directly measured *k*_obs_.

### Computational analysis of in vivo versus in vitro kinetics of eIF2α^P^ de-phosphorylation

Following inactivation of the cognate eIF2α kinase the eIF2α^P^ signal in immunoblot of CHO-K1 cell lysates decays exponentially with an observed rate constant (*k*_obs_) of 0.0013/sec (Chambers *et al*., 2015) to 0.0039/sec (Crespillo-Casado *et al*., 2017). Modelling the time-dependent change in the eIF2α^P^ signal (i.e., the dephosphorylation process) to a first order decay is justified by the observation that the concentration of eIF2α in CHO-K1 cells, ∼1 µM, is well below the *K*_M_ of the holophosphatase for its substrate, > 20 µM (Chen *et al*., 2015; Michaelis *et al*, 2011). Working at the catalytic efficiency of the PP1A/PPP1R15A/G-actin holoenzyme observed here in vitro (*k*_cat_/*K*_M_ ∼0.4*10^6^/sec*M), a similar enzyme present in the cell at a concentration of 4-10 nM could account for the *K*_obs_ of eIF2α^P^ dephosphorylation.

Cellular concentrations of PP1A vary by tissue (70 nM - 3 µM) (Kim *et al*, 2014), but at the median concentration (435 nM) recruitment of merely 2.5% of that pool into a PPP1R15A-containing complexes would account for the observed rate of eIF2α^P^ dephosphorylation in cells.

These crude estimates suggest that at reasonable concentrations, an enzyme comprised of the conserved C-terminal portion of PPP1R15A *could* account for the activity observed in cells.

### FACS (ISR quantitation)

As the effector protein in these assays is a PPP1R15A::mCherry fusion, the intensity of the fluorescence in the red channel (the Y axis of the 2D FAScan) reports on the level of effector protein in each cell and facilitates gating on populations expressing similar level of effector in comparing the median and standard error of the CHOP::GFP signal (ISR channel).

The ISR attenuation factor of wildtype and mutant PPP1R15A proteins (used in Fig. 3, 4 & 6D) was derived as follows:

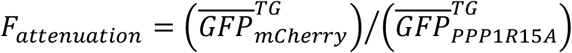

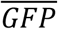, mean of the median GFP fluorescence in replicate experiments. *TG*, Thapsigargin treated cells. *UT*, untreated cells. *mCherry*, expressing mCherry alone. *PPP1R15A*, expressing the cognate wildtype or mutant PPP1R15A:: mCherry fusion protein.

Because it does not take into account the effect of the expressed PPP1R15A on the basal level of the ISR marker, this method of calculation underestimates the attenuation of the ISR especially by the more potent PPP1R15A derivatives, but as this tends to underestimate differences in the effect of various PPP1R15A derivatives on the ISR it strengthens the significance of such differences when detected.

### Quantification of stress granules

Micrographs were imported into NIH-Fuji and transfected cells expressing both mCherry and G3BP-GFP were marked as double positive and counted using the cell counter plug in. These same cells were then assigned to stress granule positive (2 or more granules) or negative (no detectable granules) marked and counted. The counts for each micrograph were exported into excel and the percent of stress granule positive cells over the total number of double positive transfected cells was calculated. 8-16 images containing an average of greater than 50 transfected cells were counted for each condition. A representative of three experiments is shown.

## DATA AVAILABILITY

The atomic coordinates and structure factors of the crystal structure of DNase I/G-actin/PPP1R15A^582-621^ complex have been deposited to the PDB with accession code PDB 7NXV. Electron microscope density maps and atomic models of the pre-dephosphorylation complex have been deposited in the EMDB and PDB, respectively, with accession codes EMD-12665 and PDB 7NZM. Plasmids, cell lines and reagents used in this study are listed in Table 3.

**Table 3.** Description of chemicals, cell lines and plasmids used in this study

| REAGENT or RESOURCE | SOURCE | IDENTIFIER |
| --- | --- | --- |
| **Bacterial strains & Cell lines** |  |  |
| T7 Express lysY/lq | New England Biolabs | Cat. No. C3013 |
| CHO-K1 with CHOP::GFP reporter | PMID:11381086 | C30 |
| **Chemicals, peptides, and recombinant proteins** |  |  |
| Thapsigargin | Calbiochem | 586005-1MG |
| Sodium arsenite | Sigma | S7400 |
| PhosTag reagent | Generon | F4002-10mg |
| Latrunculin B | AbCam | #ab14491 |
| Partially purified (DP grade) bovine pancreatic DNase I | Worthington | #LS002139 |
| Lipofectamine LTX with Plus reagent | ThermoFisher | A12621 |
| Streptavidin-coated BLI Probes | Fortebio | 18-5020 |
| **Recombinant DNA (first mentioned in the indicated figure)** |  |  |
| UK2514_Smt3_huPPP1R15A_582-621_pSUMO3 (Fig. 1b) | here | UK2514 |
| UK2699_hR15A_553-624_MBP_rPP1A_7-300_pSUMO3 (Fig. 2a) | here | UK2699 |
| UK2750_hR15A_553-624-R595E-MBP_rPP1A_7-300_pSUMO3 (Fig. 2a) | here | UK2750 |
| UK2751_hR15A_553-624-F592A_R595A-MBP_rPP1A_7-300_pSUMO3 (Fig. 2a) | here | UK2751 |
| UK2752_hR15A_553-624-R612A-MBP_rPP1A_7-300_pSUMO3 (Fig. 2a) | here | UK2752 |
| UK2755_hR15A_553-624-F592A_R595E-MBP_rPP1A_7-300_pSUMO3 (Fig. 2a) | here | UK2755 |
| UK2731_helF2a_2-187_WT_pSUMO3 (Fig. 2a) | PMID:33220178 | UK2731 |
| UK168_PerK KD-pGEX4T-1 (Fig. 2a) | PMID:9930704 | UK168 |
| UK2773_hR15A_583-621_WT_AviTag_H6_pET-30a(+) (Fig. 2dD) | here | UK2773 |
| UK2776_hR15A_583-621_R595E_AviTag_H6_pET-30a(+) (Fig. 2d) | here | UK2776 |
| UK2779_hR15A_583-621_F592A_R595E_AviTag_H6_pET-30a(+) (Fig. 2d) | here | UK2779 |
| UK2777_hR15A_583-621_F592A_R595A_AviTag_H6_pET-30a(+)<br>(Fig. 2d) | here | UK2777 |
| UK1250_FLAG_mR15A_1-653_mCherry (Fig. 3) | PMID:25774600 | UK1250 |
| UK1302_FLAG_mR15A_1-653_F585A_mCherry (Fig. 3) | PMID:25774600 | UK1302 |
| UK2724_FLAG_mR15A_1-653_F585A_R588E_mCherry (Fig 3a) | here | UK2724 |
| UK2727_FLAG_mR15A_1-653_R588E_mCherry (Fig 3a) | here | UK2727 |
| UK2775_FLAG_mR15A_1-653_F585A_R588A_mCherry (Fig 3a) | here | UK2775 |
| UK2788_FLAG_mR15A_1-653_R605A_mCherry (Fig 3a) | here | UK2788 |
| UK2830_p-eGFP-G3BP (Fig 3c) | PMID:31171631 | UK2830 |
| UK1300_FLAG_mR15A_1-653_W575A_mCherry (Fig 4) | PMID:25774600 | UK1300 |
| UK2739_pCEFL_BFP | Here | UK2739 |
| UK2740_hbActin_2-376_A204E_P243K_pCEFL_BFP (Fig 4) | here | UK2740 |
| UK2741_hbActin_2-376_A204E_P243K_E336R_pCEFL_BFP (Fig 4) | here | UK2741 |
| UK2720_hR15A_553-624-MBP_rPP1A_7-300_D64A_pSUMO3 (Fig 5) | here | UK2720 |
| UK2814_hR15A_553-624_R591A_MBP_rPP1A_7-300_pSUMO3 (Fig 6a) | here | UK2814 |
| UK2815_hR15A_553-624_R594A_MBP_rPP1A_7-300_pSUMO3 (Fig 6a) | here | UK2815 |
| UK2826_hR15A_553-624_W582A_MBP_rPP1A_7-300_pSUMO3 (Fig 6a) | here | UK2826 |
| UK2827_hR15A_553-624_R591A_R594A_MBP_rPP1A_7-300_pSUMO3 (Fig 6a) | here | UK2827 |
| UK2808_FLAG_mR15A_1-653_R584A_mCherry (Fig 6d) | here | UK2808 |
| UK2809_FLAG_mR15A_1-653_R587A_mCherry (Fig 6d) | here | UK2809 |
| UK2733_helF2a_2-187_WT_AviTag_H6_pET-30a(+)<br>(Fig 6f) | PMID:33220178 | UK2733 |
| UK2840_hR15A_553-624_W582A_MBP_rPP1A_7-300_D64A_pSUMO3 (Fig 6f) | here | UK2840 |
| UK2841_hR15A_553-624_R591A_MBP_rPP1A_7-300_D64A_pSUMO3 (Fig 6f) | here | UK2841 |
| UK2849_helF2a_2-187_M44A_pSUMO3 (Fig 7b) | here | UK2849 |
| UK2850_helF2a_2-187_K79A_pSUMO3 (Fig 7b) | here | UK2850 |
| UK2851_helF2a_2-187_Y81A_pSUMO3 (Fig 7b) | here | UK2851 |
| UK2852_helF2a_2-187_D83A_pSUMO3 (Fig 7c) | here | UK2852 |

## ACKNOWLEDGMENTS

We thank Matthieu Bollen (KU Leuven) for suggesting the PP1A^D64A^ trap, Veronica Kane Dickson, Stefan Marciniak, Joe Chambers, Alisa F. Zyryanova and Lisa Neidhardt (CIMR), Ana Crespillo-Casado (MRC-LMB) and Yao Wong (Calico) for critical comments and advice. Diamond Light Source i03 (mx21426) for X-ray crystal structure data collection. Steven Hardwick, Dima Chirgadze and Lee Cooper (Cryo-EM facility, Department of Biochemistry, University of Cambridge) for help with Cryo-EM data collection and processing. The CIMR core flow cytometry facility for help with cell sorting. The Huntington lab (CIMR) for access to the BLI Octet machine. Supported by a Wellcome Trust Principal Research Fellowship to D.R. (Wellcome 200848/Z/16/Z) and Calico Life Sciences LLC.

## AUTHOR CONTRIBUTIONS

DR, YY and HPH conceived the project, designed the experiments, analysed the data prepared figures and tables and co-wrote the manuscript. YY expressed and purified the proteins used here and conducted all the structural work, HPH conducted all cell-based and some binding experiments, DR did most of the enzymology.

## SUPPLEMENTARY FIGURE LEGENDS

**Supplementary figure 1 (related to Fig. 1).**
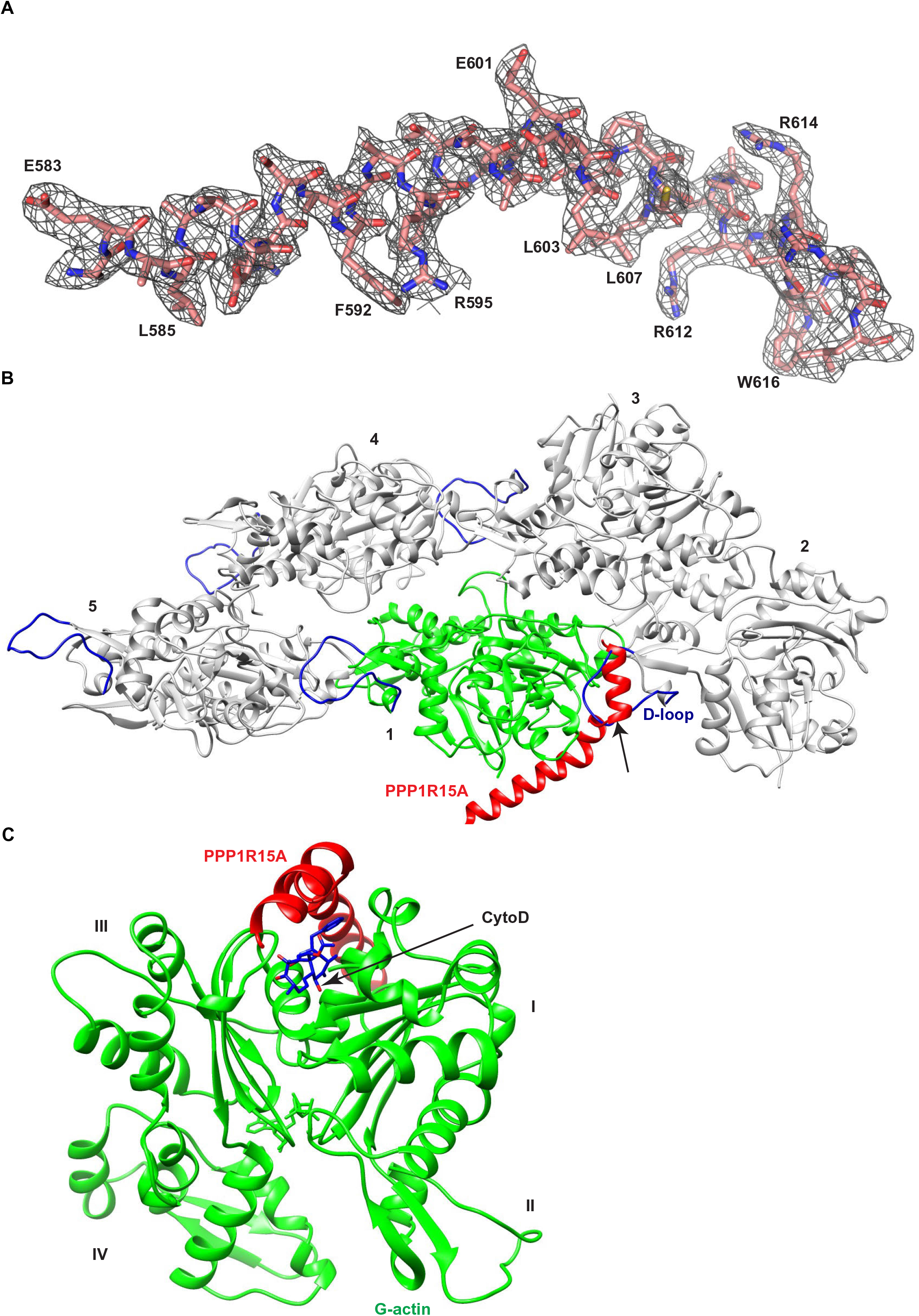
(A) Stick diagram of PPP1R15A and the corresponding 2Fo-Fc map, contoured at 1.0σ within 2.0 Å of PPP1R15A atoms. (B) Ribbon diagram of F-actin (PDB 3J8A, tropomyosin omitted) in grey with protomers numbered. The PPP1R15A/G-actin complex (in red and green respectively, with DNase I removed) has been aligned with protomer 1. Actin D-loops are shown in blue and the clash between D-loop insertion and PPP1R15A binding to G-actin is indicated by the arrow. (C) The PPP1R15A/G-actin complex with actin domains numbered and a docked molecule of cytochalasin D (from PDB 3EKS), clashing with the C-terminal helix of PPP1R15A.

**Supplementary figure 2 (related to Fig. 5).**
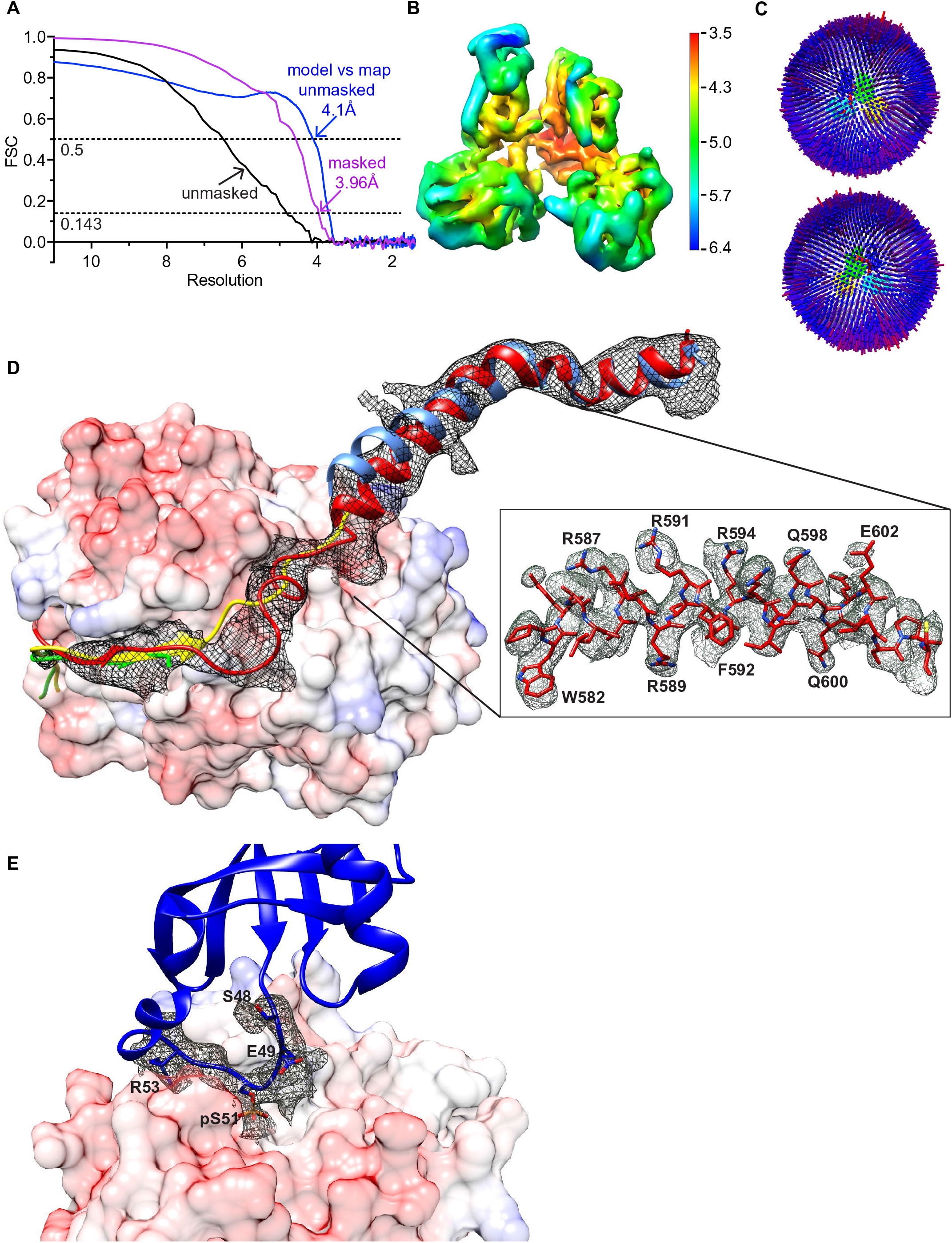
(A) The Fourier shell correlation (FSC) curves for masked (pink, produced using CryoSPARC), unmasked (black, produced using CryoSPARC) maps, and the curve for the unmasked model and map correlation (blue, produced in Phenix.real_space_refine) are shown. The resolutions at which FSC for masked map drops below 0.143 and model map correlation drops below 0.5 are indicated. (B) The 3D density map from non-uniform refinement in CryoSPARC is depicted by local resolution on a colour scale shown on the right. (C) Angular distribution plots of particles that contributed to the final map are viewed from the front and the back of the density map. The number of particles with respective orientations are represented by length and coloured cylinders (from blue to red). In the centre, the Cryo-EM map of the pre-dephosphorylation complex is coloured and angled the same as in Fig. 5A. (D) Superposition of the GS-FSC density map for PPP1R15A in the Cryo-EM pre-dephosphorylation complex (red), PPP1R15A from the crystal structure of its binary complex with PP1A (PDB 4XPN, green) and PPP1R15B from the crystal structure of its binary complex with PP1G (PDB 4V0X, yellow) (both aligned by PP1c) and PPP1R15A from the crystal structure of its binary complex with G-actin/DNase I (here, light blue, aligned by its G-actin). Inset is the density modified map of PPP1R15A^581-606^ from the Cryo-EM pre-dephosphorylation complex. (E) Density modified map for the eIF2α^P^ substrate loop (residues 48-53) on the surface of PP1A, from the Cryo-EM structure (eIF2α Arg52 sidechain was not modelled as there is no corresponding density and eIF2α Leu50 sidechain is not shown for clarity).

Supplementary movie 1

(related to Fig. 5)

Shown is the morphing of PPP1R15/G-actin between the substrate binding pre-dephosphorylation complex and the holoenzyme in absence of substrate. The holophosphatase complex (PDB 4V0U, in ivory and grey) and pre-dephosphorylation complex (in red and green) were superimposed by their PP1c moieties. The end point of the movie shows an ∼18° outward rotation of PPP1R15B/G-actin in the holophosphatase (in absence of substrate). The arrow indicates a clash that would occur with the substrate in the pre-dephosphorylation complex (blue).

